# miRNA modules for precise, tunable control of gene expression

**DOI:** 10.1101/2024.03.12.583048

**Authors:** Rongrong Du, Michael J. Flynn, Karan Mahe, Monique Honsa, Bo Gu, Dongyang Li, Sean E. McGeary, Viviana Gradinaru, Ralf Jungmann, Michael B. Elowitz

**Affiliations:** Division of Biology and Biological Engineering, California Institute of Technology, Pasadena, CA 91125, USA; Howard Hughes Medical Institute, California Institute of Technology, Pasadena, CA 91125, USA; Faculty of Physics and Center for Nanoscience, Ludwig Maximilian University, 80539 Munich, Germany; Max Planck Institute of Biochemistry, 82152 Martinsried, Germany; Department of Systems Biology, Harvard Medical School, Boston, MA 02115, USA

## Abstract

Accurate control of transgene expression is important for research and therapy but challenging to achieve in most settings. miRNA-based regulatory circuits can be incorporated within transgenes for improved control. However, the design principles, performance limits, and applications of these circuits in research and biotechnology have not been systematically determined. Here, combining modeling and experiments, we introduce miRNA-based circuit modules, termed DIMMERs, that establish precise, tunable control of transgene expression across diverse cell types to facilitate imaging, editing, and gene therapy. The circuits use multivalent miRNA regulatory interactions to achieve nearly uniform, tunable, protein expression over two orders of magnitude variation in gene dosage. They function across diverse cell types, and can be multiplexed for independent regulation of multiple genes. DIMMERs reduce off-target CRISPR base editing, improve single-molecule imaging, and allow live tracking of AAV-delivered transgene expression in mouse cortical neurons. DIMMERs thus enable accurate regulation for research and biotechnology applications.

## INTRODUCTION

Biomedical research and biotechnology heavily rely on transgene expression in living cells. The ability to accurately establish transgene expression at desired levels is critically needed in many contexts. For example, in gene therapy, overexpression of therapeutic transgenes can be toxic^1^. Similarly, in gene editing and imaging applications, overexpression can reduce specificity or increase background, respectively. However, popular expression systems, including DNA transfection and AAV vectors, as well as integrating systems such as lentivirus^2^ or piggyBac transposons^3^, typically generate a broad range of expression levels, due to variability in the number of gene copies taken up, integrated, and expressed by each cell, as well as gene expression noise^4,5^. Selecting individual stable clones can reduce variability but is time-consuming and impossible for gene therapy.

What is needed is a simple gene regulation system that could compensate for unavoidable variation in delivery and expression (**Figure 1A**). The ideal system should have several key features: First, it should be genetically compact for compatibility with most delivery vectors. Second, it should allow predictive tuning of the expression setpoint. Third, it should permit construction of multiple independent (orthogonal) regulation systems for simultaneous control of multiple genes. Finally, it should operate across multiple cell types (portability) (**Figure 1B**).

**Figure 1.**
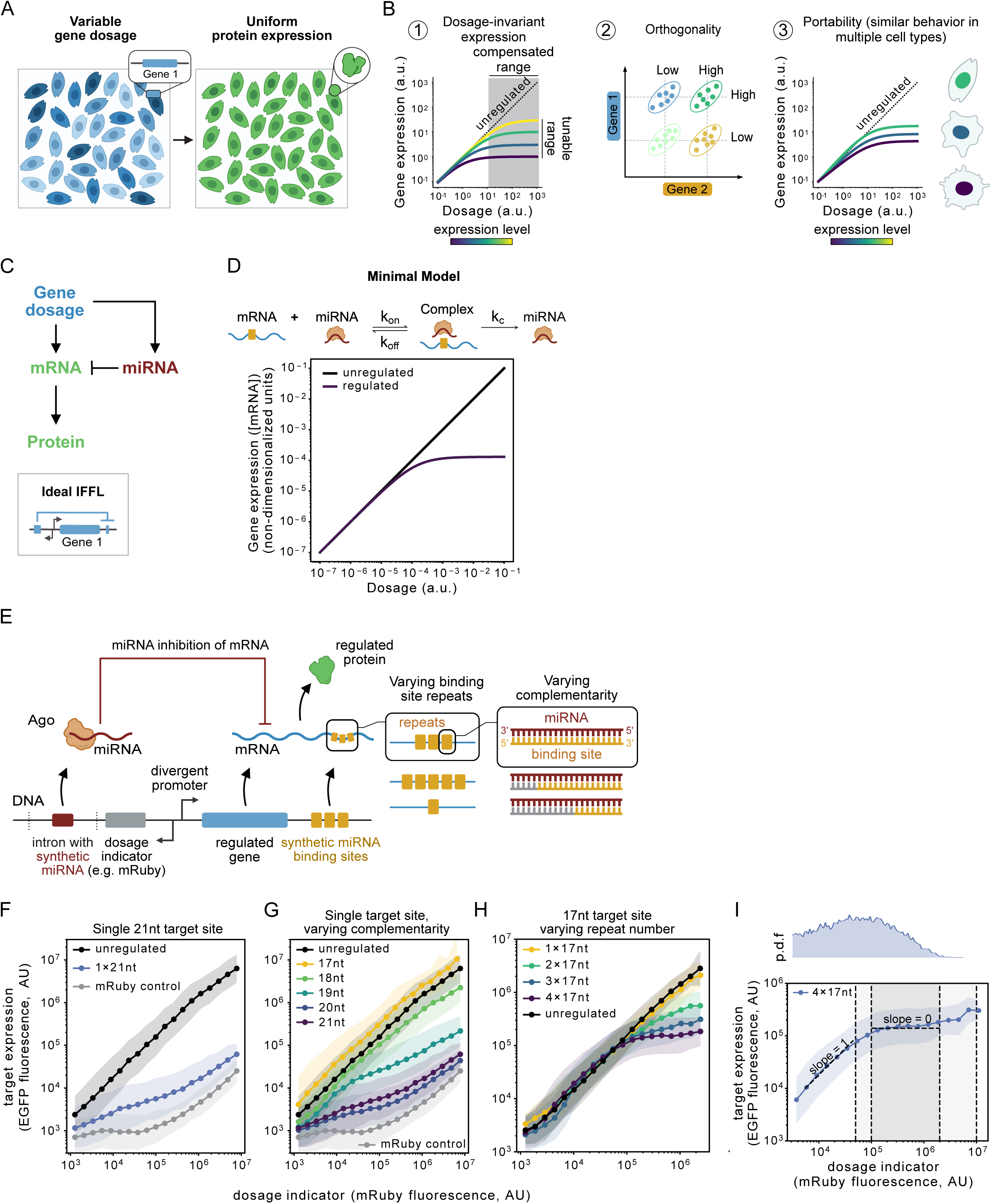
miRNA incoherent feedforward circuits enable dosage-invariant gene expression. **(A)** An ideal gene expression system generates uniform protein expression (right) levels despite variable gene dosage delivered (left). **(B)** An ideal system would enable tunable (⍰), orthogonal (⍰) control of the target and operate in multiple cell contexts (⍰). (⍰) shows that two genes can be regulated by two orthogonal circuit designs with tunable expressions. Ellipses, the majority of protein expression profiles. Dashed lines, mean expressions. **(C)** The architecture of the incoherent feedforward loop (IFFL), top panel. Simplified miRNA-based IFFL circuit configuration, bottom panel. **(D)** The minimal model shows that miRNA-mediated regulation enables dosage compensation compared to the unregulated control. Upper panel, miRNA-mediated regulation reactions. See also **Data S2**. **(E)** The circuit configuration of the miRNA-mediated IFFL. **(F)** A single, fully complementary miR-L site (1×21nt) shows strong repression compared to the unregulated but does not achieve dosage compensation. The mRuby control shows the bleed-through signal from mRuby3 in the EGFP channel. Cells are gated and binned by mRuby3 intensities. Each dot corresponds to the geometric mean fluorescence intensity of mRuby3 bin breaks and median fluorescence intensity of EGFP in the bin. Shaded regions, geometric standard deviation. **(G)** Reducing the complementarity (pairing length starting from the seed region) of the single-site target weakens the repression but does not achieve dosage compensation. **(H)** Multimerizing the 17nt site restores the repression and achieves dosage compensation at four repeats. **(I)** A zoom-in of the 4_X_17nt in **(H)**. p.d.f, probability distribution function of mRuby3 fluorescence intensity. The gray and light gray rectangles indicate the dosage range where the fluorescence intensity of EGFP does not change or change by 4-fold, respectively. The dashed lines with slope=0 or slope=1 indicate the linear dosage dependence at low dosage or dosage independence at high dosage, respectively. **(F)-(H)**, the x-axis and the y-axis show the dosage indicator expression and the target expression, respectively, both in arbitrary units (a.u.).

The incoherent feed-forward loop (IFFL) circuit motif provides an ideal foundation for these capabilities^6^. A circuit in which a target gene and its negative regulator are both encoded in the same DNA construct represents an IFFL-like configuration in which gene dosage, considered as an input, modulates expression of both the target and its negative regulator. When these effects effectively cancel out, target expression can asymptotically approach a constant expression level independent of gene dosage (**Figure 1C**).

Previous studies have introduced synthetic circuits based on this principle. Bleris et al. showed that an IFFL based on microRNA (miRNA hereafter) as the negative regulator could achieve dosage compensation^7^. Strovas et al. introduced a similar design incorporating a natural miRNA and multiple repeats of its binding site within the target gene, and examined its expression dynamics over several days^8^, achieving dosage compensation over a ∼20-fold range at the cost of potential crosstalk with endogenous genes. Yang et al later introduced an “equalizer” architecture that combined transcriptional negative feedback through the TetR protein with feed-forward miRNA regulation^9^. This generated an extended regime of strong dosage compensation but required expression of a bacterial protein. Tradeoffs between circuit complexity and efficiency are worth exploring further. Finally, while this work was under review, Love et al demonstrated other configurations of miRNA-based IFFL circuits for dosage compensation^10^. Nevertheless, fundamental questions have remained unclear: What sequence and circuit design principles optimize dosage compensation? Can these systems allow expression tuning, multiplexing of independent regulatory systems in the same cell, and portability across cell types and delivery modalities? And can they provide durable control in vivo for gene therapy applications?

To address these goals, we combined mathematical modeling, synthetic design, and quantitative circuit analysis to create a set of miRNA-based dosage compensation systems termed DIMMERs (Dosage Invariant miRNA-Mediated Expression Regulators). These circuits take advantage of multivalent miRNA regulation through the natural TNRC6 scaffold system^11^. They allow systematic tuning of expression levels by modulating the number of miRNA cassettes, numbers of target binding sites, and miRNA-target site complementarity. Further, they can be used to orthogonally regulate multiple genes in the same cell, and operate similarly across different cell types. A toolkit of ten mutually orthogonal ready-to-use expression systems can be incorporated into diverse systems. They facilitate biological imaging, improve CRISPR base editing, and function in vivo to allow AAV gene therapy applications. DIMMERs should thus allow routine research and biotechnology applications to operate with greater precision, control, and predictability.

## RESULTS

### A minimal model shows that dosage compensation requires linear sensitivity to miRNA

To guide the design of DIMMERs, we first developed a minimal model of a miRNA-based IFFL circuit. This model makes several assumptions: (1) Primary miRNA (pri-miRNA) and target mRNA are each transcribed constitutively at a fixed ratio of rates, and in direct proportion to gene dosage. (2) There is a constant total rate, per gene copy, of RISC complex production, reflecting the combined process of pri-miRNA transcription, post-transcriptional processing, and binding to Argonaute proteins^12^. (3) RISC and its target mRNA bind reversibly to form a RISC-mRNA complex. Finally, (4) formation of this complex leads to degradation of the bound mRNA.

In certain regimes, this model exhibits dosage-invariant expression profiles, in which target protein expression levels increase linearly and then asymptotically approach a dosage-independent limiting expression level (**Figure 1D**). Several parameters modulate the limiting expression level while preserving asymptotic dosage compensation. These include the binding and unbinding rates of mRNA to miRNA, and the catalytic rate of mRNA degradation. Accessing the dosage compensation regime requires that mRNA levels be linearly sensitive to RISC concentration. Below, a more detailed model in which the total amounts of free and bound mRNA were considered, revealed that miRNA-dependent catalytic degradation rates must exceed a minimal value for the total mRNA to show dosage invariance (**Data S1**). Together, these results suggest conditions in which miRNA-based IFFL circuits could produce gene dosage-invariant expression.

### Multimerization of weak target sites provides dosage compensation

Based on these results, we designed an initial set of regulatory circuits. Briefly, a synthetic miRNA and a target mRNA were placed in opposite orientations relative to a central divergent promoter^7^ (**Figure 1E**). In one orientation, a previously characterized synthetic miRNA (miR-L, based on a Renilla luciferase sequence^13^) was co-expressed with the fluorescent protein mRuby3, serving as a dosage indicator. The miRNA expression cassette included the miR-E backbone for pri-miRNA expression^13^ and was incorporated within a synthetic intron^14^. In the opposite orientation, we inserted a constitutively expressed EGFP target gene, with a single fully complementary 21-nt miRNA target site in the 3’-UTR. This format allowed analysis of multiple miRNA and target site configurations, and independent modulation of miRNA and target gene expression levels. We also systematically analyzed a broad variety of other architectures (**Data S2**).

To quantitatively measure the behavior of the circuit, we transfected U2OS cells with each circuit construct and analyzed expression by flow cytometry 48h later. We then plotted target EGFP expression versus gene dosage, indicated by mRuby3 fluorescence (**Figure 1F**). Compared to an unregulated control with no miRNA target site, the circuit strongly reduced target EGFP expression by 1–2 orders of magnitude. However, it failed to achieve dosage compensation (**Figure 1F**). We also analyzed similar circuits in which the complementary region was systematically reduced in single nucleotide increments from 21nt to 17nt. Constructs with 19 or 20nt sites retained repression but failed to produce dosage compensation, while shorter sequences lost repression altogether (**Figure 1G**). Thus, single target sites with varying levels of complementarity did not provide dosage compensation.

Native miRNAs are known to use much shorter complementary regions, including central mismatches (bulges)^12^, and multivalent interactions mediated by TNRC6 scaffold proteins^11,15^. Therefore, we next considered designs with reduced complementarity and multimerized target binding sites. Tandem repeats of two to four copies of the 17nt target site, which was inactive in a single copy, progressively increased regulation (**Figure 1H**). More importantly, they successfully reduced dosage sensitivity, particularly at higher expression levels (**Figure 1H, I**). Thus, 4 tandem 17nt sites yielded only a ∼4-fold change in expression over a ∼200-fold range of dosages (**Figure 1I**). We also tested tandem repeats of two to four copies of the 18nt and 19nt target binding sites. Interestingly, the 18nt target site showed dosage compensation behavior at 3 copies, while the 19nt target site exhibited dosage compensation with only 2 copies (**Figure S1A, B**). “Bulged” target sites that provided no regulation individually nevertheless exhibited strong regulation and limited dosage compensation when multimerized (**Figure S1C, D**). These results indicate that multiple tandem copies of individually weak target sites can achieve dosage compensation over broad dosage regimes, at varying setpoints.

### TNRC6 and Ago2 play key roles in regulation of multimerized 17nt targets

Why do multimerized weak sites produce better dosage compensation than stronger individual target sites? We reasoned that the TNRC6 scaffold protein could facilitate inhibition of multimerized weak target sites. To test whether regulation of multimeric weak sites requires TNRC6, we expressed T6B, a previously identified TNRC6B protein fragment that competitively inhibits TNRC6 activity (**Figure 2A**)^16^. T6B abolished regulation by a co-transfected 4×17nt DIMMER (**Figure 2B**), but had little effect on the single fully complementary 21nt construct (**Figure 2C**). As expected, negative controls using a T6B variant lacking the Ago2-binding domain failed to abolish regulation (**Figure S1E**). Interestingly, overexpression of wild-type Ago2 exhibited similar effects as ectopic T6B expression, nearly eliminating regulation in the 4×17nt case (**Figure 2D**), without affecting the 1×21nt configuration (**Figure 2E**), possibly due to sequestration of TNRC6. Thus, TNRC6 activity is required for regulation of the multimeric 4×17nt target, but not the single fully complementary site.

**Figure 2.**
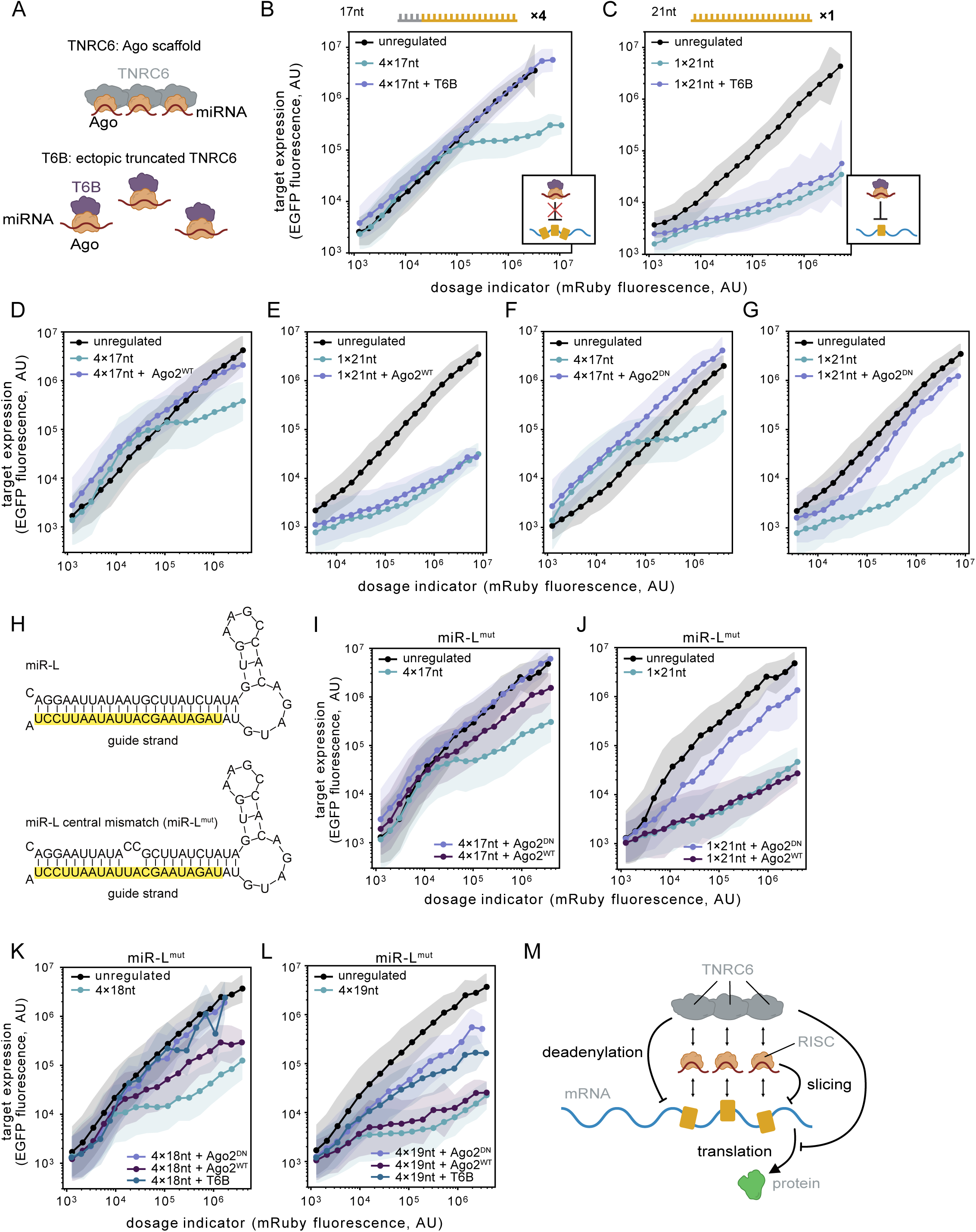
TNRC6 and Ago2 play key roles in regulation of multimerized weak targets. **(A)** T6B peptide competitively inhibits TNRC6-dependent regulation (schematic). **(B, C)** T6B expression suppresses miR-L-mediated regulation of 4×17nt construct **(B)** but not 1×21nt **(C)** construct. **(D, E)** Ectopic wild type Ago2 (Ago2^WT^) suppresses inhibition of 4×17nt **(D)** but not 1×21nt **(E)** construct. **(F, G)** Dominant negative Ago2 (Ago2^DN^) suppresses regulation of both 4×17nt **(F)** and 1×21nt **(G)** constructs. **(H)** Design of synthetic miR-L cassettes without (top) or with (bottom) central mismatches in the passenger strand. **(I, J)** Ectopic Ago2^WT^ or Ago2^DN^ modulate miR-L^mut^-mediated regulation of 4×17nt **(I)** and 1×21nt **(J)** constructs. **(K, L)** The effect of T6B/Ago2^WT^/Ago2^DN^ overexpression in the miR-L^mut^-mediated regulation for 4×18nt **(K)** and 4×19nt **(L)** constructs. **(M)** TNRC6 and Ago2 play important roles in dosage compensation (schematic). In **(B)-(C)**, **(D)-(G)**, and **(I)-(L)**, in unperturbed groups, cells were co-transfected with filler plasmids to match the total transfection amount. See also **Figure S1**.

Although Ago2 predominantly directs repression through a slicing-independent mechanism, slicing occurs in at least a dozen complementary targets^17^, and is required for maturation of two erythroid miRNAs that undergo non-canonical biogenesis^18^. To understand whether slicing activity is required for dosage compensation, we ectopically expressed a dominant negative Ago2 mutant (D669A) lacking slicing activity. This perturbation reduced regulation of both fully complementary and multimerized partially complementary targets (**Figure 2F,G**), suggesting that Ago2 slicer activity is required for repression of both targets. Further, to test for a role in cleavage of passenger strands, we incorporated a mismatch at 10-11nt in the original miRNA design (**Figure 2H**) in order to eliminate the requirement for Ago2 cleavage in miRNA maturation^18^. Inclusion of the mismatch slightly enhanced regulation of both the 4×17nt and 1×21nt (**Figure S1K, L**), possibly by elevating the efficiency of miRNA biogenesis. Although passenger strand removal is not required in this case, dominant negative Ago2 expression nevertheless largely eliminated regulation of both targets, consistent with a role for Ago2-dependent slicing in target repression (**Figure 2I-J**). By contrast, wild-type Ago2 overexpression had a partial impact on the 4×17nt target but no impact on the 1×21nt target (**Figure 2I-J**). We also analyzed other target sequences. The 4×18nt and 4×19nt targets responded similarly to T6B and Ago2^DN^, with Ago2^DN^ showing a slightly stronger de-repression effect. By contrast, wild-type Ago2 had weaker or no suppressive effect on these stronger target sites (**Figure 2K-L**), possibly because their longer complementary regions were less sensitive to indirect sequestration of TNRC6. Taken together, these results suggest that TNRC6 is required for repression of multimerized partially complementary targets but not fully complementary targets, and that Ago2-dependent slicing is required for repression of partially and fully complementary targets.

Taken together, these results suggest a potential explanation for why dosage compensation requires multimerized weak binding sites and TNRC6 (**Figure 2M**). Briefly, fully complementary binding sites are known to produce much more efficient slicing than partially complementary sites^19^. The fully complementary 21nt target could therefore produce dosage compensation at dosages and setpoints comparable to or lower than detection limits in the flow cytometry experiments. At the same time, the higher affinity of fully complementary binding sites would shift the “tail” of elevated expression due to bound-but-not-yet-degraded complexes to lower dosages. These combined effects could together explain the shape of the single 21nt fully complementary construct (**Figure 1F**). By contrast, constructs with multimerized weak binding sites (e.g. 4×17nt) would engage at higher gene dosages due to the lower intrinsic RNA affinity. This would lead to elevated setpoints and shift the “tail” of non-compensated expression to correspondingly higher dosages (**Figure 1I, Data S1**). Efficient mRNA degradation could be achieved by enhancing the intrinsically weaker Ago2-dependent slicing activity through stabilization of Ago2-mRNA complexes by TNRC6 and through TNRC6-dependent deadenylation, as described previously.

### Dosage-invariant expression levels can be tuned

Having demonstrated dosage invariant circuit designs and gained insight into their design from the model, we next sought to identify experimentally tunable parameters that modulate expression level setpoints (**Figure 1E**). We first examined varying the length of target complementarity from 8 to 21nt, while maintaining 4 tandem repeats (**Figure 3A, Data S2**). Repression was modest at 8nt, diminished with increased complementarity in the central region, and then strengthened again as more complementarity was added after the central region (**Figure 3A**, left panel). These results are consistent with previous observations that miRNA inhibition does not increase monotonically with complementarity^20,21,22^. For the miR-L target site, repression was most sensitive at 16-20 nucleotides of complementarity (**Figure 3A**). Three designs—4×17, 4×18, and 4×19— achieved dosage invariant expression, but did so at distinct setpoints spanning more than an order of magnitude in saturating expression level (**Figure 3B**). These results indicate that site length can tune setpoint.

**Figure 3.**
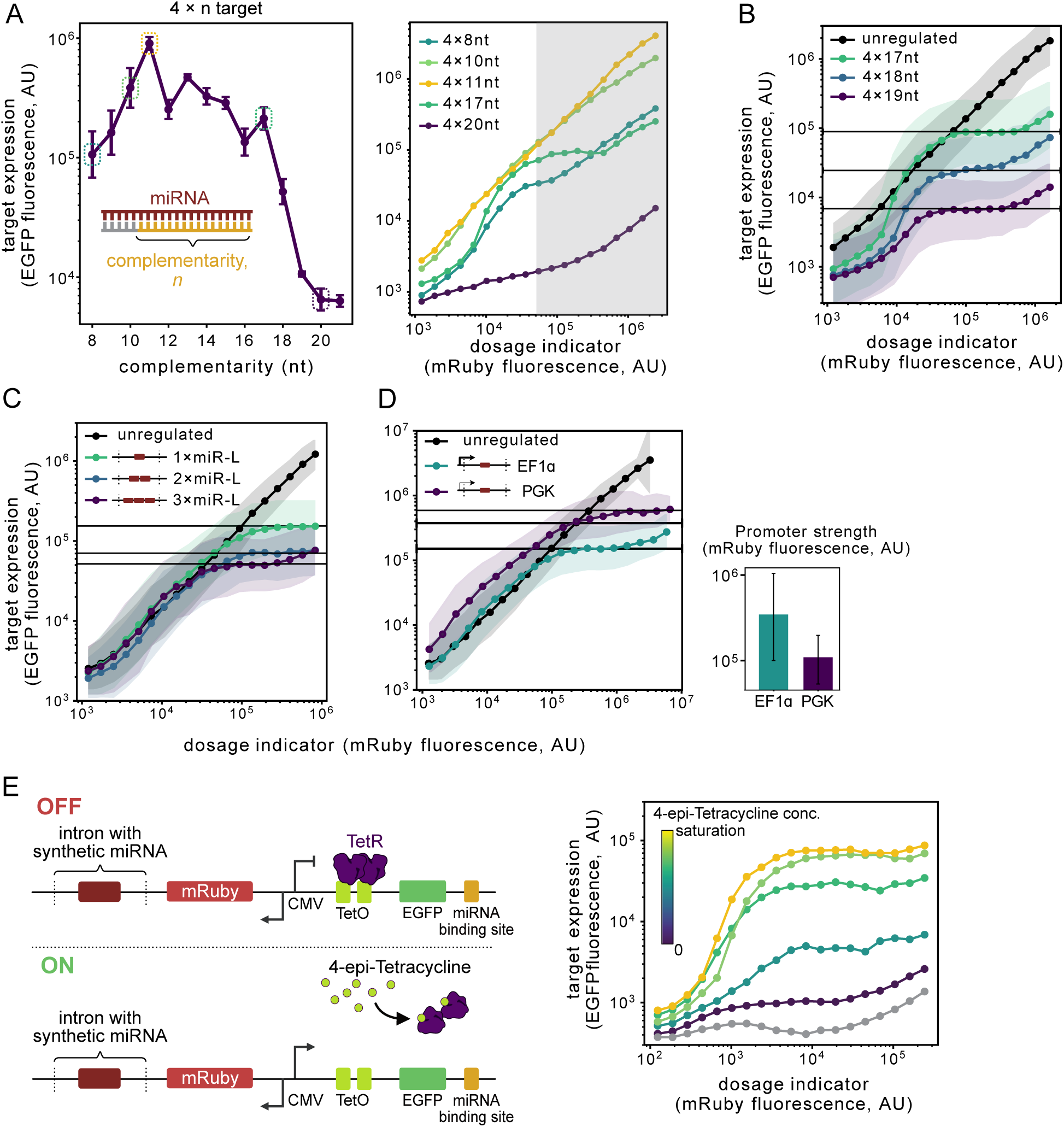
Dosage-invariant expression levels can be tuned. **(A)** 4×miR-L target expression levels exhibit non-monotonic complementarity dependence. Expression (y-axis) represents median EGFP fluorescence intensity in cells with mRuby3 fluorescence intensity > 5×10^4^, see also the shaded region in the right panel). Data represents 3 biological replicates. Right panel, dosage response curves of selected constructs (dashed boxes in the left panel). **(B)** Complementarity of 4×17-19nt monotonically regulates setpoint (black horizontal line). **(C)** Number of miR-L cassettes fine-tunes setpoint (black horizontal line). **(D)** miR-L promoter strength regulates setpoint (black horizontal line). Bars indicate mRuby3 median fluorescence intensities of transfected cells. Data represents 3 biological replicates. In C and D, the target is 4×17nt. **(E)** Tet system allows tunable dosage compensation. Left panel, design of the inducible DIMMER. A CMV promoter harboring two downstream TetO sites drives 4-epi-Tc inducible expression in the TRex cell line. Right panel, the inducible 4×19nt DIMMER dosage response curve. Drug concentrations from purple to yellow were 0, 10, 33.3, 100, 333.3 ng/mL. Gray curve, mRuby-only transfection control. See also **Data S2**.

We next varied the number of copies of the miRNA expression cassettes in the synthetic intron, effectively modulating the stoichiometric ratio of miRNA to mRNA (**Figure 3C**). Compared to a single copy, two or three copies of the miRNA reduced expression by 2-fold and 3-fold, respectively, while preserving dosage compensation, providing a means of fine-tuning expression control.

We also compared different promoters to vary transcription of the miRNA and target cassettes. For the miRNA the weaker PGK promoter allowed ∼3.5-fold more target gene expression at a given dosage level compared to the stronger EF1α, but nevertheless preserved dosage compensation (**Figure 3D**). Finally, we incorporated a tetracycline-responsive system (**Figure 3E**). This allowed setpoint tuning over ∼2 orders of magnitude while preserving dosage compensation (**Figure 3E, Data S2**). Taken together, these results demonstrate that dosage invariance can be preserved while allowing multiple mechanisms of coarse (site length, target transcription rate) and fine (miRNA promoter and copy number) setpoint tuning.

### Orthogonal dosage compensation circuits allow independent control of target genes

Engineered genetic systems increasingly require multiple genes and transcripts, provoking a need for multiple independent DIMMER systems based on orthogonal synthetic miRNA-target site pairs, which we term *synmiRs*. To design synmiRs, we started by generating five random miRNA sequences, labeled synmiR1-5. Each sequence contained A at position 1 in the miRNA, a cognate U at the 3’ end of a single target site and 25% GC content, similar to the structure of miR-L. We used an “open loop” system to analyze their behavior, allowing independent control of miRNA expression and measurement of its effect on a target miRNA reporter gene (**Figure 4A**). For a single fully complementary miR-L site, inhibition increased with miRNA dosage (**Figure 4B**), consistent with the earlier closed loop results (**Figure 1F**). synmiRs 1,4, and 5 repressed by at least an order of magnitude relative to a control lacking the miRNA (**Figure 4D, Data S2**). By contrast, synmiRs 2 and 3 achieved weaker repression, possibly due to subsequences containing two or more A/T pairs in the extensive region, which could destabilize the miRNA^23,24^ (**Figure S2A**). Consistent with this hypothesis, an A to G substitution at position 20 in synmiR-2 or at position 19 in synmiR-3 restored miRNA inhibition of target gene expression (**Figure 4D, Data S2**).

**Figure 4.**
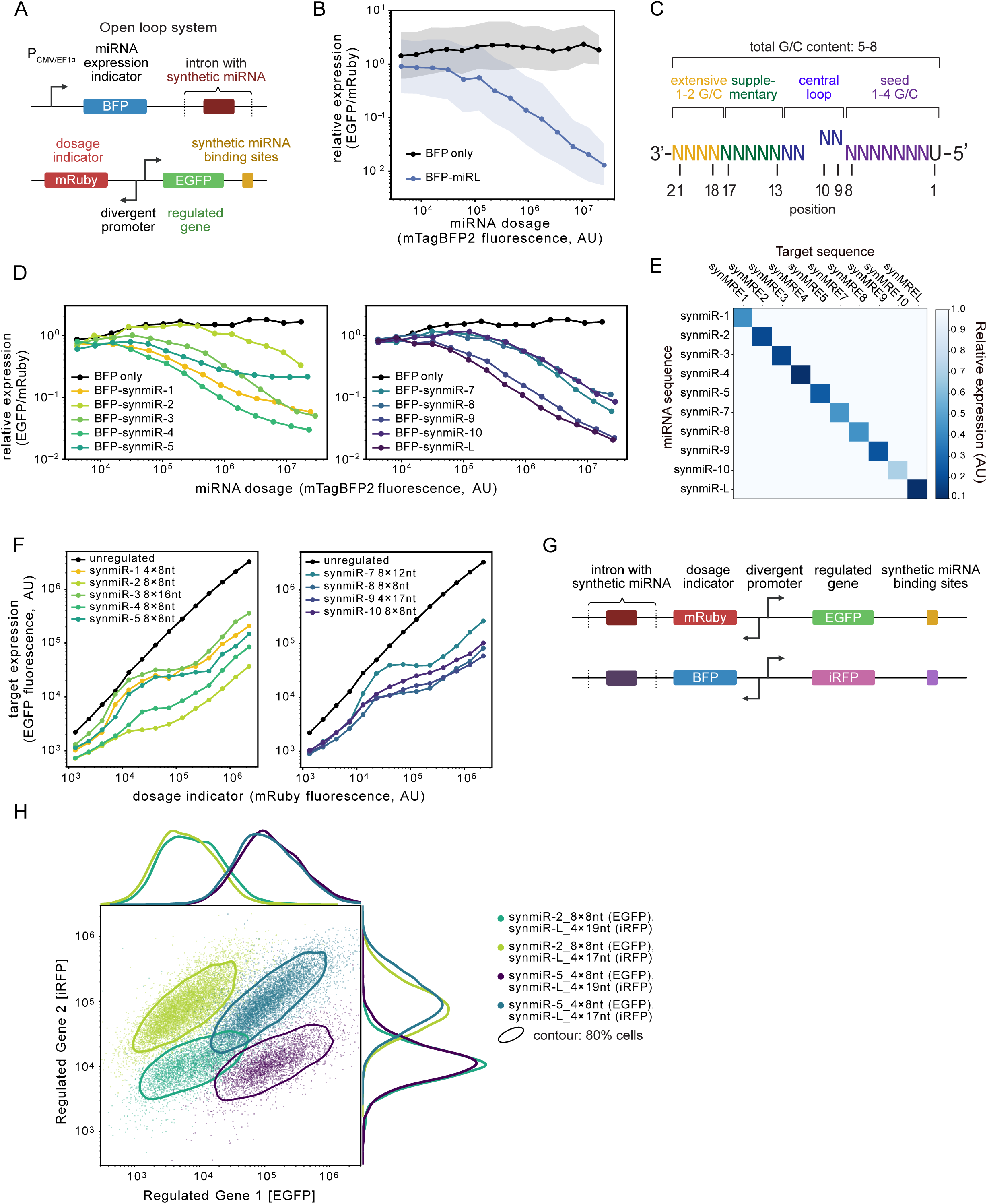
Orthogonal dosage compensation circuits allow independent control of target genes. **(A)** Open loop system allows simultaneous readout of miRNA expression (mTagBFP2) and its effect on target gene expression (EGFP), at different target gene dosages (mRuby3). **(B)** Open loop circuits reveal quantitative miRNA input-output relationship. Cells were co-transfected with the 1×21nt target, either with the miR-L construct shown in **(A)**, or mTagBFP2 as a control. Relative expression, median ratio of EGFP fluorescence intensity to mRuby3 fluorescence intensity in each mTagBFP2 bin. Dots and shaded regions are as described in Figure 1F. **(C)** synmiRs use a simple empirical design algorithm. **(D)** All synmiRs can regulate cognate single 21nt targets. **(E)** Ten synmiRs function orthogonally. Cells were co-transfected with each combination of target and miRNA for open loop analysis. Relative expression levels were quantified as described in **(B)** and normalized by maximal expression of each construct across synmiRs. **(F)** synmiRs can generate dosage compensated regulation. **(G)** Two DIMMER constructs can be analyzed in the same cells using distinct fluorescent proteins (schematic). **(H)** The double DIMMER system allows independent regulation of two genes in four distinct expression configurations. Cells were poly-transfected with the double DIMMER reporter system shown in **(G)**. Data were gated by mRuby3 and mTagBFP2 intensities. Solid contour lines indicate regions containing 80% of cells. Profile plots show the distributions of the corresponding fluorescent proteins. See also **Figure S2-S3**, and **Data S2**.

Based on these results, we formulated an empirical algorithm for synmiR design. Briefly, we generate random 21nt candidate miRNA sequences in which the mature miRNA (1) includes a 5’-U, based on known requirements for miRNA loading^12,25^, and, (2) includes 5–8 G or C nucleotides, with 1–4 of them in the seed region, and 1–2 in the extensive region (**Figure 4C**). We synthesized five candidate sequences (synmiRs 6-10) based on this simple algorithm and analyzed their open loop behavior (**Figure 4D**, **Figure S2B**). All five sequences generated strong repression as single sites, comparable to that of miR-L. Altogether, these results produced ten miRNA sequences capable of strong repression in their fully complementary form (**Figure 4D, Data S2**). Pairing each of these ten miRNAs with all ten of the target sequences in the open loop system revealed strong orthogonality in regulation, as desired (**Figure 4E, Figure S2C**).

Using these synmiR sequences, we developed a set of ten orthogonal dosage compensation circuits, using the framework in **Figure 1E**. Each of these sequences could produce dosage compensation in some configurations and dosage regimes. For example, with synmiR-4 and synmiR-5, the 4×17nt configuration produced a strong inhibition profile more similar to the fully complementary target (**Figure S2D**). We reasoned that this could reflect higher GC content in the seed and supplementary regions of these two miRNAs compared to miR-L, allowing shorter 8 or 9 nt target sites, present in 4-8 repeats, to produce dosage compensating designs (**Figure 4F**). In a similar way, we identified dosage compensation regimes for the other sequences (**Figure 4F**). Eventually, we identified nine DIMMER circuits exhibiting different levels of dosage compensation, of which five showed at least a 30-fold dosage-invariant range (**Figure 4F, Data S2**). Further, expression levels in these circuits were sensitive to the number and complementarity of target sites, allowing setpoint tuning (**Figure S3A**). These designs relied TNRC6 for dosage compensation (**Figure S3B**). The exception was synmiR-2 8×8nt, which maintained strong repression even under the T6B perturbation (**Figure S3B**). Taken together, these results provide a toolkit of dosage compensating systems and, more generally, suggest that it should be relatively feasible to engineer many additional dosage compensating systems with varying expression setpoints.

With multiple dosage compensation systems, it should be possible to independently specify the expression of multiple target genes in the same cells (**Figure 1B, panel 2**). To test this possibility, we constructed a second set of dosage compensation expression systems using distinct fluorescent reporters (**Figure 4G**). We transfected cells with pairs of systems that had different regulatory setpoints, and analyzed the resulting expression profiles of the two regulated target genes (**Figure 4H, Figure S2E**). Altogether, we analyzed four pairs of systems. Each produced a distinct two-dimensional expression distribution based on the setpoints for the two reporters (**Figure 4H**). By contrast, the unregulated group showed higher setpoints, and broader distributions of both reporters. The dosage indicators also exhibited the same distributions among all groups. Interestingly, the circuits allowed precise control of the stoichiometry of the regulated proteins (**Figure S2E**). The engineered dosage compensation systems thus make it possible to specify two-dimensional expression distributions, and suggest that control of higher dimensional distributions of more genes should also be accessible.

### Dosage compensation systems are portable and minimally perturbative

An ideal dosage compensation system would be portable, able to operate similarly across different cell types, function in both transient transfection and genomic integration, and minimally perturb the host cell. To examine these features, we transiently transfected several circuit variants, including the 4×17nt miR-L system (**Figure 1H**), in four mammalian cell lines: U2OS^26^, CHO-K1^27^, HEK293^28^, and N2A^29^. In each cell line, we observed strong and qualitatively similar dosage compensation (**Figure 5A**). Cell lines varied in the threshold dosage at which expression saturated (**Figure 5A**, gray vertical line), and in the saturating expression level (**Figure 5A**, gray horizontal line), as measured in arbitrary fluorescence units. However, the ratio of these values was conserved (**Figure 5B**). We obtained similar results for other circuits as well, including synmiR-4, with 8 repeats of a 9nt target site, as well as both synmiR-L and synmiR-5, each with 8 repeats of an 8nt target site (**Figure S4A**). Again, the ratio of the saturating expression level to the threshold dosage was similar, for each construct, across cell lines (**Figure S4B**). This suggests a model in which the miRNA circuit functions equivalently in different cell types, but protein expression strengths vary, possibly due to differences in translational capacity or basal protein degradation rates^30–33^. Together, these results indicate that the dosage compensation circuits can function across different cell contexts.

**Figure 5.**
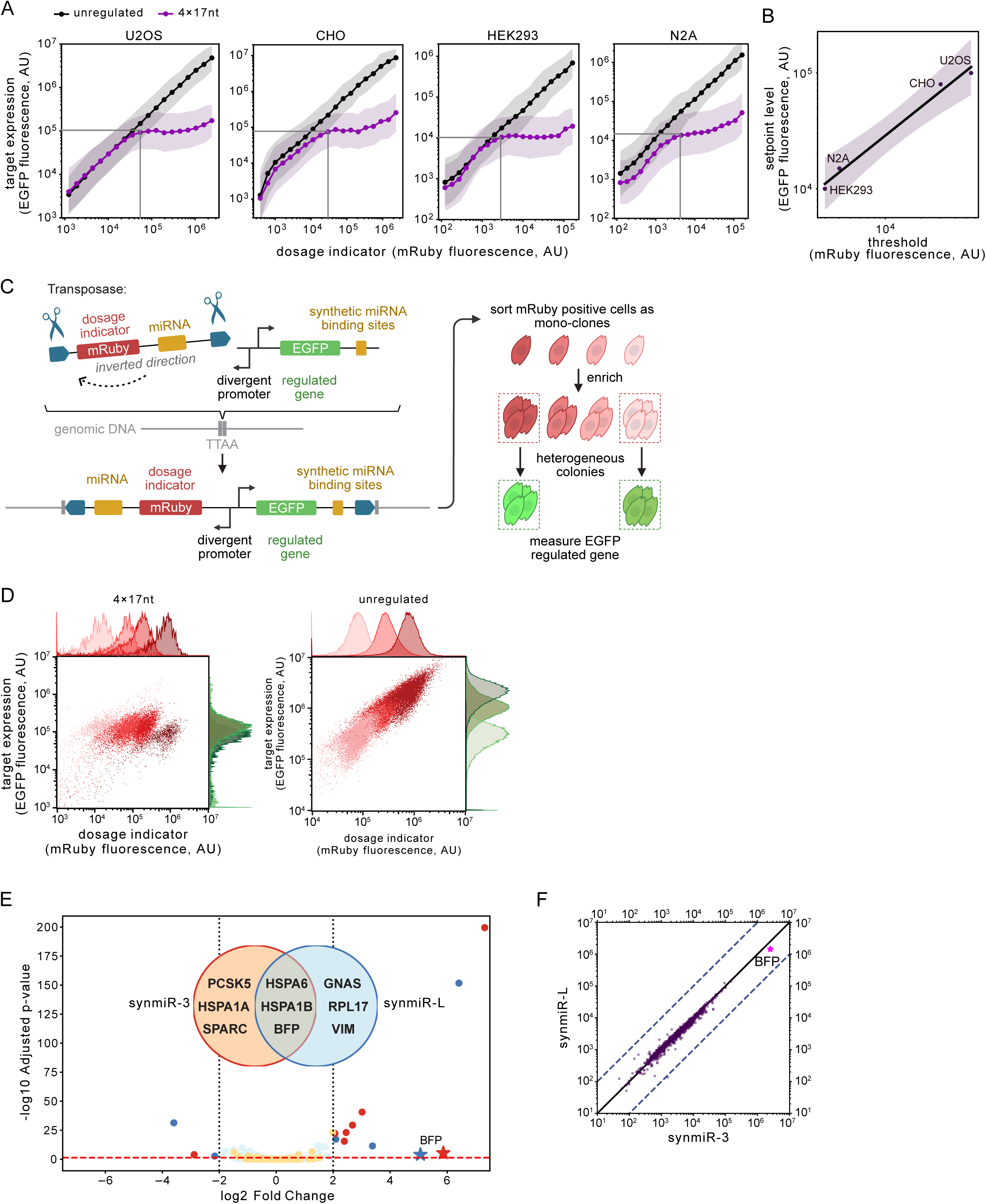
Dosage compensation systems are portable and minimally perturbative. **(A)** DIMMERs function in multiple cell lines. Gray vertical lines, minimal dosage for the compensation regime (threshold); horizontal lines, setpoint level. **(B)** The threshold and the setpoint level co-vary across cell lines. Black solid line, data linear fit in the logarithm space (**STAR Methods**). **(C)** iON transposition genomically integrates DIMMER circuits. **(D)** DIMMER circuits enable uniform protein expression (y-axis) across a range of stable integrations (x-axis). Each color shows the 4×17nt DIMMER (left panel) compared with the unregulated construct (right panel) in PiggyBac-integrated monoclonal U2OS cells. Profile histograms show mRuby3 and EGFP distributions of each mono-clone. **(E)** DIMMER circuits show limited perturbation to the endogenous transcriptome. Bulk RNAseq volcano plots show cells transfected with either synmiR-3 (orange and red dots) or synmiR-L (light blue and dark blue dots), compared to mTagBFP2-only transfected cells. Vertical dashed lines, | log_2_ (fold change) |= 2 . Horizontal red dashed line, adjusted p-value=0.05. The Venn diagram shows differentially expressed genes between synmiR-3 and synmiR-L. **(F)** Global mRNA expression levels in normalized transcripts per million (TPM) of miR-L expressing cells are plotted against those of synmiR-3 expressing cells. Solid line, equal expression in both samples. Dashed lines, 10-fold expression differences. See also **Figure S4** and **Data S2**.

Stable cell lines are important in research as well as applications like cell therapy. To find out whether dosage compensation circuits could also function in a stable integration context, we used PiggyBac transposition together with the iON system that allows expression only from constructs that have successfully integrated in the genome and undergone site-specific recombination^34^. We then selected mono-clones, and analyzed reporter expression by flow cytometry (**Figure 5C**). Integration copy numbers varied among clones by over two orders of magnitude, as indicated by mRuby3 fluorescence intensity (**Figure 5D left panel**, x-axis). Nevertheless, the cargo EGFP expression remained nearly constant (**Figure 5D left panel,** y-axis). By contrast, the unregulated mono-clones exhibited an apparent correlation between the integrated copy numbers and the EGFP expression (**Figure 5D right panel**). Thus, dosage compensation circuits function in stable integration settings.

The expression of synthetic miRNAs could in principle perturb endogenous gene expression. To identify such effects, we performed bulk RNA sequencing on cells transfected with miR-L and each of the 9 orthogonal synmiRs, and compared them to a negative control transfection of a BFP expression vector. Only a few genes were significantly up- or down-regulated by the miRNA (**Figure 5E**). These were enriched for heat shock proteins such as HSPA6. Critically, the gene sets up-regulated by different miRNAs exhibited strong overlap (**Figure 5F**, **Data S2**). Thus, for the synmiRs described here, off-target regulation appears to only reflect non-specific effects of miRNA expression, rather than sequence-specific perturbations.

### Dosage compensation enhances biological imaging

Single-molecule imaging approaches such as DNA-PAINT (Point Accumulation for Imaging in Nanoscale Topography) often rely on ectopic expression of tagged proteins^35^, which can dramatically exceed endogenous expression, distorting subcellular localization patterns. DIMMER circuits could potentially address this issue by limiting ectopic protein expression.

To test this, we transfected a DIMMER-regulated and unregulated EGFR-mEGFP membrane marker fusion protein expression-constructs into CHO-K1 cells, which lack endogenous EGFR expression^36^ (**Figure 6A**). The DIMMER circuits successfully reduced expression, as measured by receptor density relative to an unregulated control, by at least 10-fold, as estimated from flow cytometry and confocal imaging (**Figure S5A-C**). To quantify this reduction more directly, we imaged transfected cells 48h post-transfection using the DNA-PAINT method, based on DNA-conjugated anti-GFP nanobodies targeting the intracellular mEGFP tag of the EGFR (**Figure 6B-D**). Circuit-regulated receptor mean densities were ∼15-47 times lower than unregulated receptors (**Figure 6E, Figure S5D**). Further, expression distributions were narrower for the regulated plasmids compared to the unregulated ones (**Figure 6E**). Critically, DIMMER reduced expression more homogeneously than possible by simply reducing the concentration of unregulated plasmids in transfections (**Figure 6F**, **Figure S5D**). DIMMER-regulated expression with the 4×17nt circuit achieved EGFR expression levels comparable to but lower than those of wild-type U2OS cells. Thus, DIMMER circuits allow homogeneous reduction in ectopic expression to endogenous levels.

**Figure 6.**
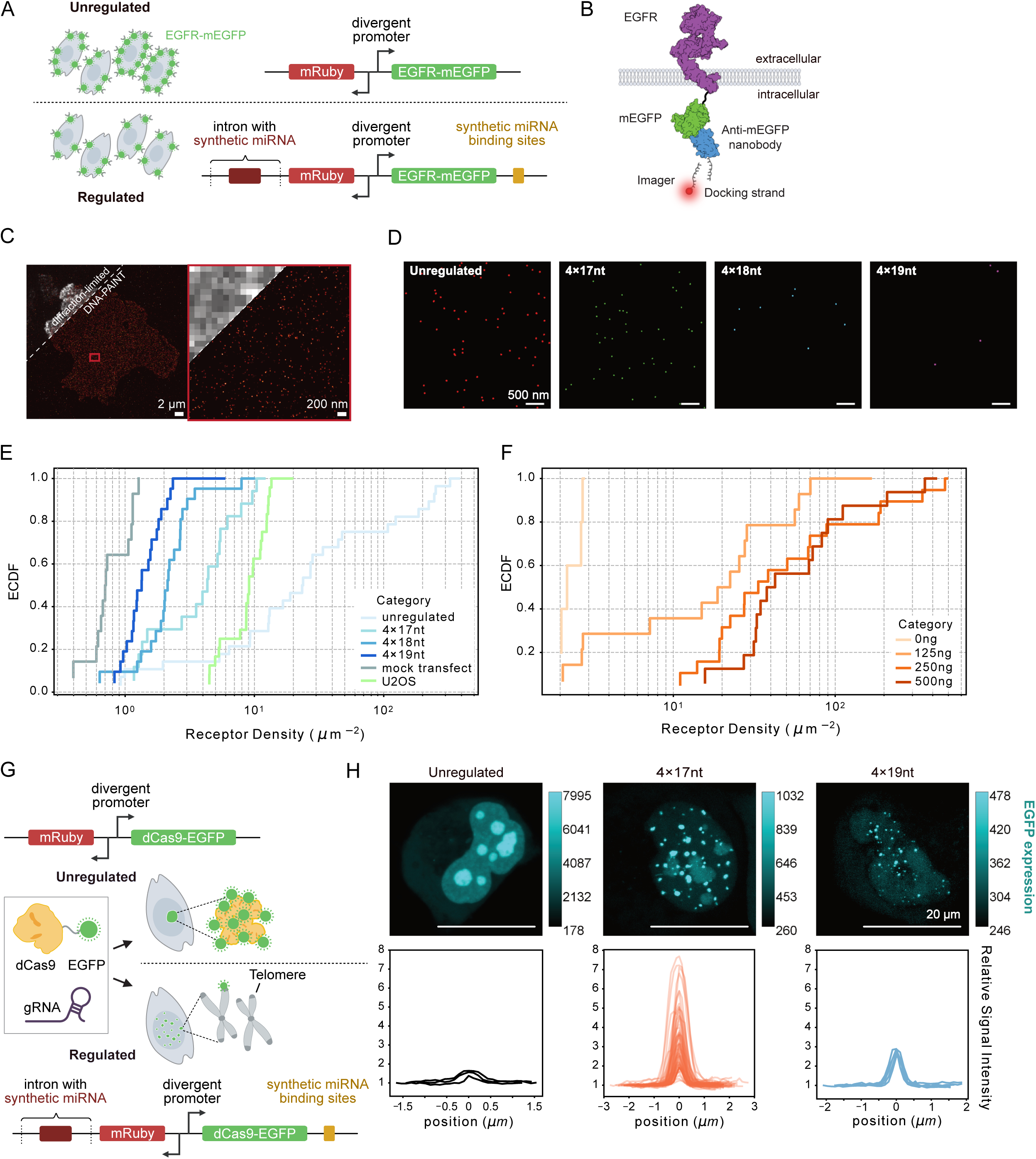
Dosage compensation circuits enhance biological imaging. **(A)** EGFR fusions to mEGFP were expressed with (lower panel) or without (upper) regulation. **(B)** EGFR (adapted from PDB 7SYE) is linked to an mEGFP labeled with a DNA-conjugated anti-GFP nanobody (PDB 6LR7). **(C)** Diffraction-limited and DNA-PAINT imaging of CHO-K1 cells expressing EGFR-mEGFP. Right panel, zoom-in of the boxed area. **(D)** Representative single-molecule images of EGFR-mEGFP. Images show (L-R) unregulated or regulated with DIMMER variants. **(E)** DIMMER circuits homogeneously reduce expression levels to near-background levels. The empirical cumulative distribution function (ECDF) of the receptor density shows CHO-K1 cells transfected with EGFR-mEGFP with or without the DIMMER module (blue curves), and the endogenous EGFR density in U2OS cells (green curve), along with mock-transfected CHO-K1 cells, measured by DNA-PAINT. Receptor densities: unregulated: 79.0 ± 42.4 µm^-2^, 4×17nt: 5.1 ± 1.7 µm^-2^, 4×18nt: 2.8 ± 1.0 µm^-2^, 4×19nt: 1.7 ± 0.5 µm^-2^, mock transfection: 0.9 ± 0.2 µm^-2^, U2OS: 9.8 ± 1.5 µm^-2^ (mean ± 95% confidence interval). **(F)** Reducing unregulated plasmid concentration does not provide a low and homogeneous receptor density, measured by DNA-PAINT. Receptor densities: 0 ng: 2.5 ± 0.4 µm^-2^, 125 ng: 36.0 ± 25.0 µm^-2^, 250 ng: 115.7 ± 74.4 µm^-2^, 500 ng: 104.1 ± 64.5 µm^-2^(mean ± 95% confidence interval). See also **Figure S5**. **(G)** dCas9 can be used to image telomeres (schematic). Telomere-targeting gRNA is driven by a U6 promoter and expressed from the dCas9-EGFP vector. **(H)** DIMMER circuits enable telomere imaging by suppressing background. Top row: representative images of dCas9–EGFP telomere labeling in the unregulated construct and in two DIMMER designs (4×17nt and 4×19nt). Color bars, measured EGFP intensities; scale bars, 20 µm. Bottom row: line-scan profiles of relative signal intensity for individual puncta. In the unregulated condition, high background obscures isolated dots. See also **Figure S6**.

CRISPR imaging methods enable analysis of specific genomic loci in cell nuclei, but can be limited by high background fluorescence due to basal expression of dCas9 fluorescent protein fusions^37,38^. DIMMER circuits also improved dCas9-EGFP imaging of telomeres. We transfected dCas9-EGFP with or without DIMMER circuits, along with a gRNA targeting repetitive telomeric sequences^38–40^ (**Figure 6G**). The circuits reduced dCas9-EGFP expression, and its dosage sensitivity (**Figure S6A**). Critically, in the unregulated system, dCas9-EGFP formed bright aggregations in the nucleolus, but seldom labeled the telomeres, consistent with previous observations^41^ (**Figure 6H left panel**). By contrast, the 4×17nt circuit restricted most fluorescence to puncta (dots), consistent with telomeric labeling^38–40^, and reduced labeling of the nucleolus (**Figure 6H middle panel**). Further, the stronger 4×19nt circuit removed nearly all labeling in the nucleolus, while maintaining apparent telomere labeling (**Figure 6H right panel**). DIMMER circuits also substantially improved signal-to-background ratio as well as contrast for individual dots (**Figure S6B**). Taken together, these results demonstrate that dosage compensation circuits can improve imaging of proteins and subcellular structures.

### DIMMER reduces off-target RNA base editing and transcriptome stress

The adenine base editor ABEMax is a powerful gene-editing tool. However, it can induce undesired transcriptome-wide off-target A-to-I editing in HEK293 cells^42^ and cause detrimental transcriptional responses^43^, raising concerns for gene therapy applications. Engineering of the TadA and/or TadA* domain has partly reduced these issues^44,45^. However, the rate of off-target RNA editing increases with the deaminase level^46^. We therefore asked whether DIMMER regulation could further reduce off-target RNA editing while preserving high levels of on-target DNA editing without additional protein engineering.

We investigated the impact of ABEMax regulation on both on-target DNA, and off-target RNA editing. To evaluate the on-target editing rate, we designed a guide RNA targeting the widely used “site 3” in HEK293^45,47^. We then co-transfected HEK293 cells with the guide RNA construct along with the ABEMax base editor, either unregulated or regulated with the DIMMER circuit module (**Figure 7A**). We initially chose the 4×18nt miR-L design since it provides an intermediate expression level setpoint. We harvested the cells 72h post-transfection, and extracted genomic DNA and RNA for further measurements.

**Figure 7.**
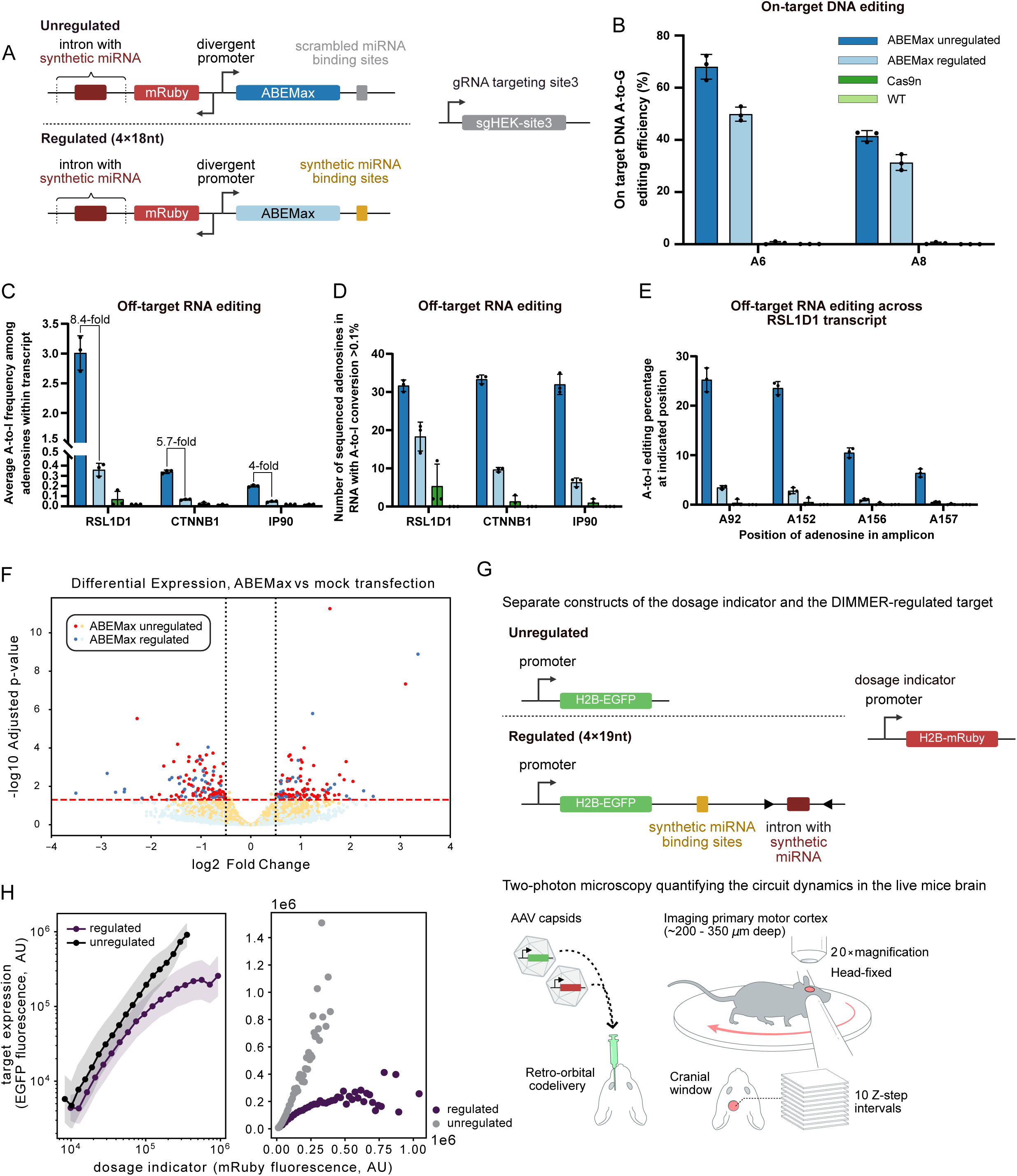
DIMMERs reduce ABEMax off-target RNA base editing and transcriptome stress, and operate in live mouse brains. **(A)** The ABEMax base editor was expressed with or without the miR-L-based DIMMER. The gRNA targeting HEK293 site 3 was expressed on a separate plasmid driven by the U6 promoter. **(B)** DIMMER modestly reduces on-target A-to-G DNA editing efficiency, measured by the percentage of the A6 and A8 edited at HEK293 site 3. **(C-E)** DIMMER strongly reduces off-target RNA editing. y-axis shows average off-target A-to-I RNA editing percentage **(C)** and the total number of sequenced adenosines in RNA with A-to-I conversion rate above 0.1% **(D)** for RSL1D1, CTNNB1, and IP90 across their transcripts, and off-target A-to-I RNA editing percentage of four specific A sites in the RSL1D1 transcript **(E)** (cf. **Data S2**). In **(B-E)**, each dot is a biological replicate. Error bars, standard deviation. **(F)** DIMMER reduces off-target transcriptome perturbation. Volcano plots show bulk RNAseq of cells containing regulated (4×18nt) and unregulated ABEMax compared to mock transfection. Black dashed lines, | log_2_ (fold change) |= 0.5 . Red dashed line, adjusted p-value=0.05. Data represents three biological replicates. **(G)** AAV-delivered constructs enable DIMMER-regulated protein expression in the mouse brain. Upper panel: the H2B-mRuby3 construct (dosage indicator) and the H2B-EGFP construct (target, either unregulated or regulated) driven by the same promoter were co-delivered to live mouse brains. Lower panel: two-photon microscopy experiments quantifying circuit dynamics in live mice brains. See details in **STAR Methods**. **(H)** In vivo brain expression of the CaMKIIα promoter cohort measured by microscope. Data were pooled, binned and plotted based on mRuby3 fluorescence intensities on logarithmic (left) or linear (right) scales. See also **Figure S7**.

The regulated ABEMax base editor performed on-target editing at a similar, if modestly decreased, rate compared to the unregulated control (**Figure 7B**). To measure the off-target RNA editing rate, we selected three transcripts (*CTNNB1*, *RSL1D1*, *IP90*) known to exhibit high off-target editing due to similarity with native TadA tRNA substrates^47, 44^. The DIMMER circuit successfully decreased the mean A-to-I editing rate by 4 to 8.4-fold across the three transcripts (**Figure 7C**). It also reduced the number of detectable A-to-I conversions within the transcript (using a cutoff of 0.1% based on the sequencing results) (**Figure 7D**). At specific adenosines that are highly edited by the unregulated ABEMax, the circuit exhibited dramatically lower (around 10-fold) editing rates (**Figure 7E, Data S2**). We did not detect off-target genomic DNA editing on the predicted potential endogenous off-target sites with or without DIMMER. These results show that DIMMER regulation can improve the ratio of on- to off-target edits.

The DIMMER circuit also reduced perturbations to the transcriptome. We performed bulk RNA sequencing on cells treated with ABEMax with or without DIMMER regulation. DIMMER reduced transcriptome changes compared with unregulated ABEMax expression (**Figure 7F, Data S2**). More specifically, unregulated ABEMax exhibited ∼2-fold more differentially regulated genes (adjusted p<0.05) compared to the regulated construct. Gene ontology annotation showed that heat shock proteins, DNA damage, and repair-associated proteins were perturbed in both unregulated and regulated ABEMax groups. However, the unregulated ABEMax group produced 89 differentially expressed genes in the ‘nucleoplasm and nucleus’ category, suggesting a more pervasive impact (**Figure 7F, Data S2, Table S2**). Together, these results indicate that DIMMER can improve the specificity of base editors by limiting their expression.

### DIMMER circuits can operate in live mouse brains

Monogenic diseases of the central nervous system could potentially be cured by gene replacement therapies^48,49^ using recently developed adeno-associated virus (AAV) capsids that deliver genes across the blood-brain barrier^50–52^. However, many therapeutic genes are toxic or deleterious when overexpressed and expression must be maintained over long periods. These considerations underscore the need for controlled, durable cell-type specific in vivo expression of transgenes.

To test whether DIMMER circuits could function in vivo in mouse brain, we designed a set of 6 AAV vectors that express H2B-EGFP with or without the broadly dosage compensating 4×19nt DIMMER circuit (**Figure 7G upper panel, Figure S7A**), For in vivo expression, we compared three promoter systems with distinct cell-type specificities: the CaMKIIα promoter (CaMKIIα hereafter), which drives expression in excitatory projection neurons, particularly in the cortex and hippocampus^53,54^; a hybrid CMV enhancer–MeCP2 promoter (MeP hereafter), which enables broad neuronal expression throughout the brain^55^; and a minimal β-globin promoter driven by the mDLX2 enhancer (mDLX2 hereafter), which selectively targets GABAergic interneurons in the forebrain, including the motor cortex^56^. As controls, we also delivered a corresponding set of unregulated constructs lacking the DIMMER circuit. All constructs were codelivered with the mRuby3 construct for internal normalization. We injected each pair of constructs into one cohort of mice, and analyzed expression of both fluorescent reporters over time by two-photon head-fixed imaging at single-cell resolution through a previously described cranial implant^57^ over the left motor cortex (**Figure 7G lower panel, STAR Methods**). This approach allowed us to monitor the fluorescent dynamics of EGFP and mRuby3 expression in individual cortical cells over timescales of 56-84 days.

To better understand the relationship between promoter strength, expression dynamics, and regulatory compensation, we analyzed circuit performance over time (**Figure S7B-E**). For all three promoters, fluorescence gradually accumulated over multiple weeks. Notably, the mDLX2 promoter, which is relatively weak and interneuron-specific, showed no evidence of dosage compensation across all timepoints (**Figure S7D**). Similarly, the MeP promoter, which drives moderate, pan-neuronal expression, shifted the distribution downward in regulated conditions but did not reach expression levels sufficient to trigger compensation, even at later time points (**Figure S7E**). In contrast, the strong CaMKIIα promoter exhibited dosage compensation at later stages (28–56 days). By 28 days, expression from the regulated CaMKIIα construct plateaued and diverged from the unregulated condition, consistent with engagement of post-transcriptional attenuation by the DIMMER circuit (**Figure 7H, S7B**). These findings suggest that dosage compensation can occur in the brain for some construct designs, that promoter strength and time-dependent expression dynamics are critical for entering the dosage compensation regime, and that weaker promoters may be unable to reach high enough expression levels to activate compensatory repression with this DIMMER variant (**Figure S7F**). In the future, increasing the number of miRNA copies or miRNA target sites could enable stronger dosage compensation with weaker promoters.

## DISCUSSION

Ectopic gene expression is a cornerstone of modern biology, and gene and cell therapy but precise control has remained elusive in most applications. The miRNA circuits described here achieve precise, sequence-tunable gene dosage-invariant control of protein expression (**Figure 2 and 3**), orthogonal control of multiple target genes (**Figure 4**), and portability across cell types and modes of delivery (**Figure 5**). We therefore anticipate that they could become standard systems for controlled gene expression in diverse areas of biomedical science and biotechnology, including imaging (**Figure 6**), CRISPR-based gene editing (**Figure 7**), and AAV-based gene therapy (**Figure 7, Figure S7**).

The process of engineering these circuits revealed unexpected design principles. Strong miRNA regulation, as obtained with the 1×21nt circuit, was not sufficient for dosage invariance within targeted expression levels (**Figure 1F**). Rather, simultaneously reducing complementarity and multimerizing target sites to engage multivalent TNRC6-dependent regulation was essential. In other regulatory systems, multispecific recognition is associated with ultrasensitivity. Here, however, it allowed linearly sensitive repression of mRNA required for dosage invariance but shifted responses to higher dosages. It will be interesting to learn whether natural miRNA regulatory systems use multispecific binding in similar ways^58,59^. We also observed that separation of miRNA and target gene into divergently transcribed genes can be useful in allowing strong, independent control of miRNA expression relative to target mRNA (**Figure 3B-E**) but is not required for precise expression, facilitating applications like gene therapy where vector capacity can be limiting (**Data S2**).

The system may be extended or improved in different ways. First, better miRNA sequence-function models could potentially allow predictive design of setpoints. Second, current expression distributions exhibit significant variability, or noise (**Figure 3E**). This could reflect transcriptional bursting of the target or the miRNA, and could also be exacerbated by potential differences in the time delays for miRNA production and processing compared to RNA splicing and nuclear export. Going forward, understanding these and other contributions to overall variability could help to reveal fundamental limits of expression precision in the cell^4,5,60,61^. Finally, the ability to combine DIMMER regulation with inducible promoters or natural enhancers for cell-type specificity could make these systems even more useful.

The circuits introduced here will be useful in diverse settings. They reduced background in imaging applications (**Figure 6**). They also reduced off-target RNA editing by CRISPR base editors, while maintaining on-target editing, suggesting they may be useful for gene editing applications (**Figure 7**). Analysis of stronger circuits with even lower expression setpoints could help to achieve even lower off-target edit rates. A major application category is gene therapy (**Figure 7G-H, Figure S7**). Many monogenic diseases that are gene therapy targets exhibit toxicity at high levels of the therapeutic gene, making it critical to suppress overexpression^62–66^. In the future, DIMMER circuits could also ensure fixed expression levels for receptors or other components in cell therapies^67^, and therefore allow expression of transcription factors and other components at physiological expression levels for regenerative medicine and other applications^68,69^. Thus, we anticipate these systems becoming useful components in a wide range of engineered research and therapeutic contexts.

### Limitations of the study

The DIMMER circuits have limitations. First, we do not yet have a predictive model of how an arbitrary miRNA sequence will behave quantitatively in a dosage compensation circuit. Varying miRNA regulation strength by modulating complementarity was straightforward with miR-L, but more complex for other synmiRs, suggesting that additional factors besides base pairing are likely important for fully predicting the activity of a miRNA on its target. Second, some aspects of the relevant molecular mechanisms remain unclear, including the role of catalytic slicer-dependent regulation^70,71^.

## RESOURCE AVAILABILITY

### Lead contact

Further information and requests for resources and reagents should be directed to and will be fulfilled by the lead contact, Michael B. Elowitz.

### Materials availability

Plasmids generated in this study are being submitted to Addgene. All unique/stable reagents generated in this study are available from the <u>lead contact</u> with a completed Materials Transfer Agreement.

### Data and code availability

- The original and processed data of this study has been deposited at CaltechDATA (data.caltech.edu) and is publicly available at the time of publication. (DOI: 10.22002/thgnh-6aj57).
- The original code to process data and mathematically model the circuits has been deposited at CaltechDATA (data.caltech.edu, DOI: 10.22002/thgnh-6aj57)
- Any additional information required to reanalyze the data reported in this paper is available from the <u>lead contact</u> upon request.

## Supporting information

List of miRNA and targets used in this study, related to Figure 1-7

Gene Ontology annotations for significantly regulated genes in ABEMax unregulated vs. mock and ABEMax regulated vs. mock, related to Figure 7

## ACKNOWLEDGMENTS

We thank Phillip Zamore (UMass), Josh Mendell (UT Southwestern Medical Center), Acacia Mayfield, James Linton, Kaiwen Luo, Martin Tran, Duncan Chadly, Shiyu Xia, Yodai Takei, Felix Horns, Lucy Chong, Leah Santat, Sheng Wang for discussion and technical support; Inna-Marie Strazhnik (Caltech) for graphical design; Fei Chen (Broad Institute), Rui Malinowski, Evan Mun, Judy Shon, Jacob Parres-Gold, and other members of the Elowitz lab for critical feedback, and administrative support. This work is supported by the National Institute Of Biomedical Imaging And Bioengineering of the National Institutes of Health under Award Number R01EB030015, and Rett Syndrome Research Trust under award number RSRT.2024. The content is solely the responsibility of the authors and does not necessarily represent the official views of the National Institutes of Health. M.B.E. is a Howard Hughes Medical Institute Investigator. We apologize for incomplete citations due to space limits.

## AUTHOR CONTRIBUTIONS

R.D., M.F., and M.B.E. conceived and designed the study. M.B.E. supervised the study. R.D., M.F., K.M., M.H., B.G., D.L., and S.E.M. performed or assisted with experiments and data analysis. R.D. and M.B.E. wrote the manuscript with input from all authors.

## STAR⍰METHODS

### EXPERIMENTAL MODEL AND STUDY PARTICIPANT DETAILS

#### CELL CULTURE

U2OS cells (ATCC Cat# HTB-96), T-Rex cells (ThermoFisher Cat# R71007), CHO cells (ATCC Cat# CCL-61), HEK293 cells (ATCC Cat# CRL-1573), and N2A cells (ATCC Cat# CCL-131) were cultured at 37 °C in a humidity-controlled chamber with 5% CO2. The growth media consisted of DMEM (Dulbecco’s Modified Eagle Medium, Thermo Fisher #11960-069) supplemented with 10% FBS, 1 U/ml penicillin, 1 μg/ml streptomycin, 1 mM sodium pyruvate, 1×NEAA (Thermo Fisher #11140-050), 1 mM L-glutamine, and 0.1 mg/mL Normocin (InvivoGen #ant-nr).

#### ANIMALS

All animals used in this study were approved by the Institutional Animal Care and Use Committee (IACUC) at the California Institute of Technology. Wildtype C57BL female mice (1.5-4 months of age) were ordered from Jackson Laboratories (Bar Harbor, ME, USA) and were used for surgeries, viral injection, and head-fixed imaging. All mice were singly-housed post-operation and for the remainder of the data acquisition in a room with a reverse light cycle (12h – 12h). For all imaging sessions, mice were imaged during the dark phase of their light cycle.

### METHOD DETAILS

#### PLASMIDS CONSTRUCTION

Constructs used in this study are listed in Key Resource Table. Some constructs were generated using standard cloning procedures. The inserts were generated using PCR or gBlock synthesis (IDT) and were ligated either by T4 ligase (NEB #M0202M) or In-Fusion (Takara #102518) assembly with backbones that are linearized using restriction digestion. *E.coli* competent cells (NEB Cat# C3040H and NEB Cat# C3019H) were utilized to amplify the plasmids. The rest of the constructs were designed by the authors and synthesized by GenScript. Selected constructs will be deposited at Addgene and the maps are available.

#### MIRNA ALIGNMENT TO THE DATABASE

Each synthetic miRNA sequence (mature miRNA, 22 nt) is aligned to the known miRNA sequence database^72–79^ (https://mirbase.org/) to identify if there are any similarities existing between the synthetic sequences and the natural sequences.

#### TRANSIENT TRANSFECTION

Cells were seeded at a density of 50,000 cells in each well of a 24 well plate (or at a density of 10,000 cells in each well of a 96 well plate), either standard for flow cytometry or glass-bottom for imaging experiments, and cultured under standard conditions overnight. The following day, the cells were transiently transfected using Fugene HD (Promega #E2311), according to the manufacturer’s protocol.

#### FLOW CYTOMETRY

Cells were incubated 2 days after transient transfection, and the culture media was replaced 24 hours post-transfection. Cells were trypsinized with 75 μL of 0.25% trypsin for 5 minutes at 37 °C. After digestion, cells were resuspended with 125 μL of HBSS containing 2.5 mg/ml BSA and 1 mM EDTA. Cells were then filtered through a 40 μm cell strainer and analyzed using a CytoFLEX S instrument (Beckman Coulter).

#### CELL SORTING

To prepare the mono-clones that expressed the genomic-integrated DIMMER circuit, cells were harvested and resuspended in sorting buffer (BD FACS Pre-Sort Buffer) supplemented with 1 U/ml DNase I by the cell sorter (Sony MA900) as mono-clones. Cells were sorted into 96 well plates in the normal U2OS culture media. Cells were expanded in the 24 well plate before flow cytometry measurement.

### BULK RNA SEQUENCING TO IDENTIFY THE OFF-TARGET EFFECTS OF THE SYNTHETIC MIRNA ON THE TRANSCRIPTOME

#### Sample preparation and sequencing

To verify the off-target effect of all the synmiRs, U2OS cells were plated on 6-well plates with 300,000 cells per well. Cells were transfected the following day with 1,000 ng of either the control plasmid or the BFP-miRNA plasmid using Fugene HD (Promega #E2311) according to the manufacturer’s instructions. Media was replaced with 2 mL of fresh media 24 hours post-transfection. Cells were harvested 48 hours post-transfection by digestion with 0.25% Trypsin-EDTA, centrifugation at 300g for 5 minutes, and removal of the supernatant by aspiration. The cell’s pellet was stored in -80 °C prior to the purification.

RNA was extracted using the RNeasy kit (Qiagen #74106) according to the manufacturer’s instructions. RNA was treated with Turbo DNase (Thermo Fisher #AM2238) and purified using the RNeasy kit RNA cleanup protocol. mRNA sequencing libraries were prepared by Novogene.

### DNA-PAINT

#### Buffers

The following buffers were used:

● NH_4_Cl solution: NH_4_Cl (Roth, no. K298.1) was dissolved in ddH_2_O for a 2 M stock solution, filtered with 0.2 μm filter.
● Blocking buffer: 1×PBS, 1□mM EDTA (Thermo Fisher, no. AM9260G), 0.02% Tween-20 (Life Science, no. P7949), 0.05% NaN_3_ (Serva, no. 30175.01), 2% BSA (Sigma-Aldrich, no. A9647-100G), 0.05□mg/ml sheared salmon sperm DNA (Life Technologies, no. 15632011), filtered with 0.2 μm filter.
● Imaging buffer: 1×PBS, 1 mM EDTA, 0.02% Tween-20, 500 mM NaCl (Thermo Fisher, no. AM9760G), supplemented with PCA, PCD, trolox, filtered with 0.2 μm filter.

#### PCA, PCD and Trolox

Trolox (100×) was prepared by dissolving 100 mg (±)-6-hydroxy-2,5,7,8-tetra-methylchromane-2-carboxylic acid (trolox; Sigma-Aldrich, 238813-5G) in 430 μL of 100% methanol (Sigma-Aldrich, 32213-2.5L), 345 μL of 1 M NaOH (VWR, 31627.290) and 3.2 mL of water.

PCD (40×) was made by mixing 154 mg of 3,4-dihydroxybenzoic acid (PCA; Sigma-Aldrich, 37580-25G-F) in 10□mL of water and NaOH and adjusting the pH to 9.0. PCD (100×) was prepared by adding 9.3□mg of protocatechuate 3,4-dioxygenase pseudomonas (PCD; Sigma-Aldrich, P8279) to 13.3 mL of buffer containing 100□mM Tris-HCl pH□8.0 (Thermo Fisher Scientific, AM9855G), 50□mM KCl (Thermo Fisher Scientific, AM9640G), 1□mM EDTA, and 50% glycerol (Sigma-Aldrich, 65516-500ml)).

#### Cloning

An mEGFP gBlock (obtained from IDT) was inserted into a pcDNA3.1(+) backbone (Thermo Fisher, no.V79020) via Gibson assembly. Two codon-optimized fragments of human EGFR (obtained from IDT) were fused to the mEGFP-pcDNA3.1(+) backbone via Gibson assembly. The plasmid concentration was measured with the NanoDrop One (Thermo Fisher Scientific).

#### Cell culture

CHO-K1 cells (ATCC: CCL-61) were cultured in Ham’s F-12K (Kaighn’s) medium (Gibco, no. 21127022) supplemented with 10% FBS (Gibco, no. 11573397). U2OS-CRISPR-Nup96-mEGFP cells (a gift from the Ries and Ellenberg laboratories) were cultured in McCoy’s 5A medium (Gibco, no. 16600082) supplemented with 10% FBS. All cells were cultured at 37 °C and 5% CO2 and split every 2-3 days via trypsinization using trypsin-EDTA (Gibco, no. 25300096).

#### Nanobody-DNA conjugation

First, the anti-GFP nanobody (clone 1H1, Nanotag Biotechnologies, N0305) and anti-rabbit IgG nanobody (Nanotag Biotechnologies, N2405) were conjugated to a DBCO-PEG4-Maleimide linker (Jena Bioscience, no. CLK-A108P). After removing the unreacted linker with Amicon centrifugal filters (10,000 MWCO), the DBCO-nanobody was conjugated via DBCO-azide click chemistry to the docking strand (Metabion, see sequence in Table S3). A detailed description of the conjugation can be found in the former work^85^.

#### Fixation of cells

The cells were fixed with 37 °C pre-warmed methanol-free 4% PFA (Thermo Fisher, no. 043368.9M) in 1×PBS for 15 min. Then, the cells were washed 3 times with 1×PBS and then permeabilized with 0.125% TritonX-100 (Sigma-Aldrich, no. 93443) in 1×PBS for 2 min. After washing 3 times with 1×PBS, the cells were blocked with the blocking buffer either overnight or for at least 3h at 4 °C.

#### Sample preparation of U2OS cells

10,000 cm^-2^ U2OS cells were seeded on an ibidi eight-well high glass-bottom chambers (no. 80807). On the next day, the cells were fixed as stated in the fixation protocol and blocked for 3h at 4 °C. After washing 3 times with 1×PBS, the primary EGFR antibody (Cell Signaling, clone D38B1, no. 4267) with a dilution of 1:200 in blocking buffer was incubated overnight at 4 °C. The next day, the sample was washed 3 times with 1×PBS and 25 nM of R2 anti-rabbit NB in the blocking buffer was incubated at RT for 1h. After washing 3 times with 1×PBS, the sample was post-fixed with 4% PFA in 1×PBS for 5 min at RT. The cells were then quenched with 200 mM NH_4_Cl (Roth, no. K298.1) for 5 min and washed 3 times with 1×PBS. 90nm gold-nanoparticles (Absource, no. G-90-100) in 1:1 in 1×PBS were incubated for 5 min at RT. After washing 3 times with 1×PBS, the cells were washed once with the imaging buffer.

#### Sample preparation for plasmid dosage measurement

5,000 cm^-2^ CHO-K1 cells were seeded on an ibidi eight-well high glass-bottom chambers (no. 80807) one day before transfection. The cells were transfected with EGFR-mEGFP plasmids with a Thermo Fisher Lipofectamine 3,000 reagent (no. L3000008) with the lower Lipofectamine concentration as indicated by the manufacturer and different plasmid concentrations (0 ng, 125 ng, 250 ng and 500 ng of plasmid per well). After 48h of transfection, the cells were fixed as indicated in the fixation protocol and blocked with blocking buffer for 3h at 4 °C. The cells were then post-fixed with 4% PFA and 0.2% glutaraldehyde (Serva, no. 23115.01) in 1×PBS for 10 min. After quenching the sample with 200 mM NH_4_Cl for 5 min and washing 3 times with 1×PBS. 90 nm gold-nanoparticles in 1:1 in 1xPBS were incubated for 5 min at RT. After washing 3 times with 1×PBS, the cells were washed once with the imaging buffer.

#### Sample preparation of DIMMER plasmids

5,000 cm^-2^ CHO-K1 cells were seeded on an ibidi eight-well high glass-bottom chambers (no. 80807) one day before transfection. The cells were transfected with EGFR-mEGFP plasmids with a Thermo Fisher Lipofectamine 3,000 reagent (no. L3000008) with the lower Lipofectamine concentration as indicated by the manufacturer and 250 ng plasmid per well (200 μL solution per well and 25 μL transfection solution). After 48h of transfection, the cells were fixed as indicated in the fixation protocol and blocked with the blocking buffer for 3h at room temperature (RT). 25 nM R3 anti-GFP nanobodies were incubated in the blocking buffer for 1h at RT. After washing 3 times with 1×PBS, the nanobodies were post-fixed with 4% PFA and 0.2% glutaraldehyde in 1×PBS for 10 min. The cells were then quenched with 200 mM NH_4_Cl for 5 min and washed 3 times with 1×PBS. 90nm gold-nanoparticles (Absource, no. G-90-100) in 1:1 in 1×PBS were incubated for 5 min at RT. After washing 3 times with 1×PBS, the cells were washed once with the imaging buffer and imaged in the imaging buffer.

#### DNA-PAINT imaging

The samples were imaged in the imaging buffer with the corresponding imager strand (obtained from Metabion, see Table S3 for imager strand sequences) for 40k frames with 100ms exposure time per frame and a readout rate of 200 MHz.

#### Microscope setup

The samples were measured on inverted total internal reflection fluorescence (TIRF) microscopes (Nikon Instruments, Eclipse Ti2) which are equipped with an oil-immersion objective (Nikon Instruments, Apo SR TIRF ×100/numerical aperture 1.49, oil) and a perfect focusing system. The mRuby3 signal was bleached by the 560 nm laser (MPB Communications, 1□W) by using Highly inclined and laminated optical sheet (HILO) illumination. Afterwards, the TIRF mode was established. The Cy3B-conjugated imagers were excited with the 560 nm laser. The laser beam was cleaned with a filter (Chroma Technology, no. ZET561/10) and coupled into the microscope with a beam splitter (Chroma Technology, no. ZT561rdc). The fluorescent signal was filtered with an emission filter (Chroma Technology, nos. ET600/50m and ET575lp) and projected onto a sCMOS camera (Hamamatsu Fusion BT) without further magnification. The camera’s central 1152×1152 pixels (576×576 pixels after binning) were used as the region of interest, with a resulting effective pixel size of 130nm. The raw microscopy data was acquired via μManager (Version 2.0.1).

### BASE EDITOR EXPERIMENT

#### Transfection and Sample collection

HEK293 cells were seeded at the density of 150k/well in the 24-well plate one day before the transfection. For each well, 750 ng ABEMax base editor plasmid and 250 ng sgRNA plasmid were co-transfected using Fugene HD (Promega #E2311), according to the manufacturer’s protocol. Cells were harvested 72 h post-transfection.

#### Amplicon sequencing

To quantify the on-target base editor editing rate, cells were sorted based on the constitutively-expressing mRuby on the base editor plasmid and EGFP on the sgRNA plasmid. Genomic DNA was extracted using Qiagen DNeasy kit (cat. nos. 69504) according to the manufacturer’s protocol. The genomic DNA amplicons were amplified using the primer in the reference^47^, with the adaptor sequence on each end. Amplicons were size-verified by DNA electrophoresis, purified by the Qiagen gel purification kit, and sequencing was performed by Genewiz Amplicon-EZ (150-500bp) service.

To quantify the off-target base editor editing rate on the genome, genomic DNA was extracted, and amplicons were obtained using the potential genomic DNA off-target loci-targeted primers reported previously^47^. To quantify the off-target editing on the RNA level, the total RNA was extracted from the cell using the Zymo Direct-zol RNA miniprep kit, according to the manufacturer’s protocol. cDNA samples were prepared using the Maxima H Minus First Strand cDNA Synthesis Kit (Thermo Fisher Scientific). Amplicons were obtained using the potential RNA off-target-loci-targeted primers reported previously^47^. The amplicons were then size-verified by DNA electrophoresis, purified with the Qiagen gel purification kit, barcoded with the Illumina Miseq 16S Metagenomic sequencing index primers, and further purified with the magnetic NGS beads (Omega Bio-Tek). Library was quantified and normalized with the Qubit fluorometer. Library was then denatured and sequenced with Element AVITI System Sequencing Instrument using AVITI 2×150 Sequencing Kit Cloudbreak (Catalog # 860-00013).

#### Bulk RNA sequencing

To analyze the transcriptome-wide perturbation of the base editor, total RNA was extracted from the cells with the Zymo Direct-zol RNA miniprep kit (lot # R2050). 50 ng of extracted mRNA from each sample were used as inputs for downstream NGS library preparation.

mRNA-seq libraries were prepared in 96-well format with a modified 3’Pool-seq protocol^90^. In brief, reverse transcription reaction were prepared by mixing input RNA with 1□μl Indexed RT Primer (10□μM), 1□μl 10□mM dNTP Mix (New England Biolabs Cat# N0447S),1□μl diluted ERCC Spike-In Mix 1 (0.004□μL stock ERCC per μg RNA, Thermo Fisher Cat# 4456740), 3.6 μl of 5×RT buffer (Thermo Fisher Cat# EP0752), 0.5 μl of RNAse inhibitor (Thermo Fisher Cat# EO0381), 1 μl Maxima RT H minus (Thermo Fisher Cat# EP0752), 2.5 μl 10 uM Template Switching Oligo into a 18 μl reaction. Reverse transcription was carried out in a thermocycler with a program described in 3’Pool-seq protocol.

Samples from each row of 96-well plate were pooled (column pooling) by mixing an equal volume of each Reverse Transcription reaction into a new well at a total volume of 20□μl. Residual primers were then degraded with the addition of 1□μl Exonuclease I (New England Biolabs) and incubated at 37□°C for 45□min followed by denaturation at 92□°C for 15□min. Subsequent cDNA amplification, tagmentation, and row pooling was performed following 3’Pool-seq protocol.

Finally, 20 μl of pooled NGS library were subject to Gel-based size selection using E-Gel EX Agarose Gel (Thermo Fisher Cat# G401001) to enrich for fragments with size range between 200-1000 bp and eluted in 15 μl.

Eluted pooled NGS libraries were examined in an Agilent TapeStation 4200 (Agilent Technologies) to determine average fragment sizes. Library concentration was quantified in a Qubit 3.0 Fluorometer (Life Technologies). NGS library molarity was then calculated using 660□g/mol per base-pair as a molecular weight. NGS libraries were diluted to 2 nM, denatured in 0.2 N NaOH, and loaded onto Element AVITI sequencer following Element Biosciences Cloudbreak Sequencing user guide.

### DNA-FISH IMAGING

To cross-validate the success of DIMMER-regulating dCas9 labeling the telomeres, Nikon 2 Multipoint acquisition was used to record the absolute coordinates of the fields of view. After the live cell imaging, the plate was removed from the scope and DNA-FISH was performed. Dye-conjugated probes designed to bind to the distal telomeric regions of the chromosome were used following the previously reported protocol^91,92^. The plate was then loaded on the scope, and the recorded coordinates were retrieved. The exactly same fields of view were matched by eye around the recorded coordinates.

### DIMMER CIRCUIT DYNAMICAL QUANTITATION IN MICE CRANIAL WINDOW

#### AAV Viral Production

All constructs were cloned using the cloning strategy described above. The viral titers of the viruses used in this study were given in Table S4. CAP-B22 and CAP-B10 are AAV capsids developed in the previous study^51^. The unit of the viral titers is viral genome copies per mL.

#### Cranial Window Surgeries

Mice were deeply anesthetized under 1-2% isoflurane and given subcutaneous injections of ketofen (5 mg/kg), buprenorphine (3.25 mg/kg), dexamethasone (2 mg/kg) and saline for analgesia and hydration. A subcutaneous injection of a few drops of bupivacaine was given under the incision site for local analgesia. An incision was performed over the skull and a piece of skin near the left motor cortex was removed to make room for the implant. A circular craniotomy of approximately 3-mm in diameter was drilled over the left motor cortex and centered 1.6-mm anterior and 1.6-mm lateral from bregma. Saline was flushed periodically throughout the drilling over the bone to hydrate and prevent heating. After extraction of the cranial bone, a gelatin sponge soaked in saline was applied over the exposed dura to remove debris and stop any residual bleeding. A small drop of Kwik-sil silicone elastomer was applied over the dura and a 3-mm diameter glass coverslip (#1.5) was implanted over the craniotomy and sealed with dental cement. A custom-made titanium headbar was fixed using cement over the window to allow for later head-fixation during 2-photon imaging. Following surgery, mice were allowed to recover for at least 4-days before beginning habituation to a rotary platter setup.

#### AAV Vectors and Administration in Mice

AAV packaging, purification, and delivery were performed as described in the earlier study^93^. Following surgical implants of cranial windows and recovery, mice were anesthetized under 1-2% isoflurane. Intravenous administration of AAVs was performed by injection into the retro-orbital sinus of mice. All vectors were each delivered at a total of 5×10^12^ viral genomes (VG) in a volume of 50 µL per mouse from 2-4 months of age. A total of three cohorts were designed to test for the performance of the DIMMER circuits with different gene-regulatory elements: CMV-enhancer paired with a minimal MeCP2 promoter (CMV enhancer-MEP229), an mDLX2 enhancer paired with a minimal beta-globin promoter (mDLX2-minBglobin), and a CaMKIIα promoter. Each cohort had two groups: One group co-expressing an unregulated H2B-EGFP transgene and an unregulated H2B-mRuby3 transgene, and a regulated group co-expressing a 4×19nt DIMMER regulated H2B-EGFP transgene and an unregulated H2B-mRuby3 transgene. These transgenes were co-delivered separately in their own AAV-capsids. The mRuby3 signal was later used as an internal reference for the EGFP signal for each mouse.

#### Two-Photon Head-Fixed Imaging and Acquisition

Mice were habituated to the handler’s hand for 3 days. Following this, they were head-fixed onto a rotary platter for an additional 3 days to further habituate to the imaging setup. After habituation, mice were imaged over a variable number of days and weeks to capture the progression of EGFP and mRuby3 expression. We used a custom home built 2-photon microscope equipped with a galvo-galvo scanner for imaging. Each imaging session involved acquiring two high-resolution Z-stacks per mouse using a 20×/1.0 NA objective (Zeiss W Plan-Apochromat, Cat. No. 421452-9700-000) with water immersion. Each of these Z-stacks consisted of 10 Z-steps, ranging from 200-250 µm to 300-350 µm below the dura, with each Z-step separated by 10 µm intervals, for a final 100 µm thick Z-stack. Each Z-step was acquired at a scan rate of 1 Hz with a resolution of 256 by 256 pixels and a pixel dwell time of 13 µs for a total of 180 seconds, over an area of 400 µm × 400 µm. Z-steps in each Z-stack were later processed as mean-intensity projected TIF files to generate a single Z-step image. Each Z-stack was first acquired at 940 nm wavelength (EGFP), followed by 1100 nm (mRuby3). The average laser power beneath the objective was maintained across imaging sessions at 36-37 mW for the 940 nm wavelength and 46-47 mW for the 1100 nm wavelength.

### QUANTIFICATION AND STATISTICAL ANALYSIS

#### FLOW CYTOMETRY DATA ANALYSIS

We used the FlowJo V10 and self-built python code to analyze the flow data. The regression curves and the confidence intervals in Figure 5B and Figure S4 were computed using the statsmodels.regression.linear_model.OLS package in logarithm spaces. The statistical details of each experiment are described in the results and figure legends.

#### IN-VITRO IMAGE ANALYSIS

The transiently transfected U2OS/HEK293 cells were imaged using a Nikon confocal microscope at 60× magnification, such that each image was spaced by 0.5 microns in the z-direction. For HEK293 cells, 1 slice in the z-direction was taken given the relatively smaller volume compared to U2OS. Images were processed by the Fiji software^80^.

To analyze the relative signal intensity in the dCas9 imaging experiment, maximum intensity projection of 11 slices of the z-stacks were applied. To determine the signal intensity of the dots, freehand lines were drawn to select the dot regions. To determine the signal intensity in the background, ∼ 5 micron-long straight lines centering the dots were drawn. The signal intensities were generated by the Fiji ROI mean intensity function. The relative signal intensity is calculated by normalizing the background intensity to be 1.

To analyze the signal to noise ratio (SNR), freehand lines were drawn to select the dot regions and the nucleus regions. The noise intensity was calculated by the intensity in the nucleus excluding the dots area. The SNR was calculated dividing the mean intensities of each dot by the noise.

### BULK RNA SEQUENCING DATA ANALYSIS TO IDENTIFY THE OFF-TARGET EFFECT OF THE SYNTHETIC MIRNAS

#### Preprocessing of sequencing data

Reads from the RNA sequencing were aligned to a custom reference genome using kallisto (0.48.0)^81^. This reference consisted of the human genome GRCh38 cDNA (https://github.com/pachterlab/kallisto-transcriptome-indices/releases) and mTagBFP2 coding sequences. Weakly expressed genes were filtered out if they exhibited fewer than 3 samples expressing at least 10 transcripts per million (TPM), or if the maximum TPM among all samples was less than 105. Then, filtered counts were input to DEseq to eliminate the impact of size factors. As the BFP-only cells were used as a reference to evaluate the off-target effect of the miRNAs, genes that showed fluctuating expressions among the three biological replicates of BFP-only cells should be removed from analysis. To achieve this, we computed log(1 + *x*) where *x* denotes the normalized TPM among the three biological replicates of BFP-only cells. The Fano factors^82,83^ of these logarithmic expressions were determined and ranked. Transcripts that ranked as the largest 2.5% in logarithmic Fano factors were eliminated from further analysis. Finally, we computed log(1 + *x*), where *x* denotes the normalized TPM among all the samples. A difference function was defined to compute the absolute value of the log(1 + *x*) difference between each sample and the untransfected sample. The medians of the difference function of the BFP-only groups and the experimental groups were calculated and used for comparison. The difference between those two difference functions were ranked and similarly, transcripts that ranked as the largest 3% were removed from further analysis.

The Fano factor is defined as:

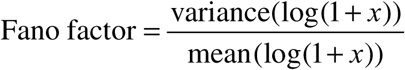

The equation of the difference function is defined as:

Δ(BFP, untransfected) =| log(1+ *x*_BFP_) – log(1 + *x*_untransfected_) |

Δ(experimental, untransfected) =| log(1+ *x*_experimental_) – log(1 + *x*_untransfected_) |

Δ_ranked_ = median(Δ(BFP, untransfected)) – median(Δ(experimental, untransfected))

#### Differential gene expression analysis

To characterize the perturbations that synthetic miRNA brought to the endogenous transcriptome, differential expression analysis was performed using DESeq2 (1.40.1)^84^ in R (4.3.1) comparing transcript counts in miRNA transfected cells and BFP-only cells.

### DNA-PAINT DATA ANALYSIS

Obtained fluorescent data was reconstructed with Picasso software^35^. The data was first drift-corrected with redundant cross-correlation, after that with picked gold particles as fiducials.

In order to determine the receptor density, a homogeneous area of the cells was picked and the DNA-PAINT data was clustered with the SMLMS clustering algorithm of Picasso^35,86^.

The identified cluster centers were used to calculate the measured receptor density per μm^2^ (number of cluster centers per area). Given that the labeling efficiency (LE) of the binders to their targets was less than 100%, a correction factor was applied to account for incomplete labeling. The EGFR density was calculated by multiplying the measured receptor density with the LE of the respective binders. The LE values of the binders were determined as previously described in the former work^87^. Specifically, the LE of the anti-GFP nanobody was determined to be 37%, while the LE of the EGFR antibody was 71%.

To test the statistical significance of the determined receptor densities, we performed the Mood’s median test. Mood’s median test is a non-parametric statistical test which tests whether the median of two groups are statistically different. Our statistical tests were performed with a custom python script using functions from the scipy.stats module^88^.

### BASE EDITOR EXPERIMENT DATA ANALYSIS

#### Amplicon sequencing data analysis

The on-target and off-target editing rates were analyzed using the online tool CRISPResso2^89^.

#### Bulk RNA sequencing data analysis

Weakly expressed genes were filtered out if they exhibited fewer than 3 samples expressing at least 10 transcripts per million (TPM). Differential expression analysis was performed using DESeq2 (1.40.1)84 in R (4.3.1). Identified significantly differentially expressed genes were eliminated from further analysis if they were identified among the unregulated, regulated, and untransfected groups with the same trend, which indicates that these genes tend to be expressed as more similar profiles among the base editor-transfected groups and the WT group, thus very likely to be the artifact of the filler-plasmid expression. GO annotation was performed using the DAVID web server (https://davidbioinformatics.nih.gov/).

### TWO PHOTON MICROSCOPY DATA ANALYSIS

Two-photon Z-stacks were acquired separately for EGFP and mRuby3 channels and processed independently. Raw data files were segmented into individual z-step movies based on predefined frame intervals using a custom MATLAB script, with each segment saved as a multi-page TIFF file^94^. Motion correction was applied to each TIFF using rigid registration, and motion-corrected data were used to generate mean- and maximum-intensity projection images for each z-step. Following motion correction, mean-intensity projection images were flatfield corrected using MATLAB’s imflatfield function with a specified radius of 20 pixels to remove uneven illumination artifacts. Corrected images were saved in a dedicated output directory for subsequent analysis. To ensure spatial correspondence between channels, each flatfield-corrected mRuby3 image was aligned to its corresponding EGFP image using normalized cross-correlation, correcting for lateral drift and maximizing overlap between channels.

Cell segmentation was performed on the mRuby3 channel using the Cellpose^95^ deep learning-based segmentation algorithm (model type = “cyto”, diameter = 9.0 pixels), generating individual cell masks for each z-step image. These masks were saved in both TIFF and NumPy formats and served as a consistent reference for fluorescence quantification across channels. For each z-step, the corresponding mRuby3-derived cell mask was applied to the EGFP image to extract the total EGFP fluorescence per cell by summing the pixel intensities within each ROI. Separately, a dedicated script was used to quantify total mRuby3 fluorescence per cell by applying the same cell mask to the original mRuby3 image for that z-step. Results were saved as per-z-step .csv files, with all individual pixel values archived in cumulative text files for downstream statistical analysis and visualization.

To evaluate single-cell expression and dynamics across time, total EGFP and mRuby3 fluorescence values per cell were aggregated across multiple mice and imaging timepoints. For the CaMKIIα and CMVe-MeCP2 cohorts, z-steps 1, 5, and 10 were selected from each 10-step Z-stack to reduce the likelihood of overcounting the same cells across adjacent planes. For the mDLX2-minβglobin cohort, only z-step 5 was selected.

## SUPPLEMENTAL INFORMATION

Document S1. Figures S1–S7, Data S1, Data S2, Tables S3 and S4

Table S1, List of miRNA and targets used in this study, related to Figure 1-7

Table S2, Gene Ontology annotations for significantly regulated genes in ABEMax unregulated vs. mock and ABEMax regulated vs. mock, related to Figure 7

