## Supplementary material for "miRNA modules for precise, tunable control of gene expression": Figures S1-S7, Data S1, Data S2, Tables S3 and S4

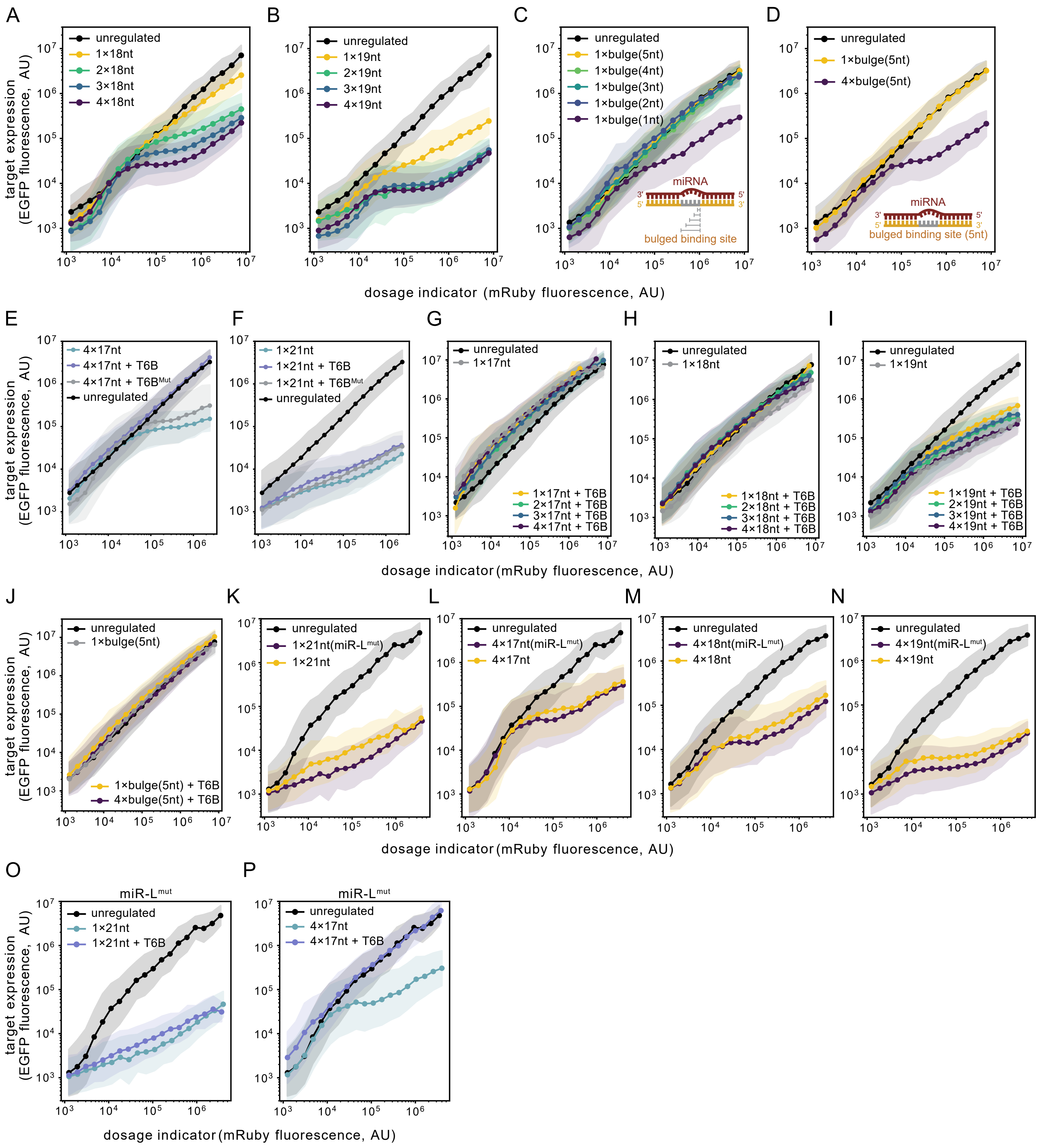
A
B
C
D
E
F
G
H
I
J
K
L
M
N
O
P
target expression (EGFP fluorescence, AU)
dosage indicator (mRuby fluorescence, AU)
unregulated
1×18nt
2×18nt
3×18nt
4×18nt
1×19nt
2×19nt
3×19nt
4×19nt
1×bulge(5nt)
1×bulge(4nt)
1×bulge(3nt)
1×bulge(2nt)
1×bulge(1nt)
4×bulge(5nt)
miRNA
bulged binding site
bulged binding site (5nt)
4×17nt
4×17nt + T6B
4×17nt + T6BMut
1×21nt
1×21nt + T6B
1×21nt + T6BMut
1×17nt
1×17nt + T6B
2×17nt + T6B
3×17nt + T6B
4×17nt + T6B
1×18nt + T6B
2×18nt + T6B
3×18nt + T6B
4×18nt + T6B
1×19nt + T6B
2×19nt + T6B
3×19nt + T6B
4×19nt + T6B
1×bulge(5nt) + T6B
4×bulge(5nt) + T6B
1×21nt(miR-Lmut)
4×17nt(miR-Lmut)
4×18nt(miR-Lmut)
4×19nt(miR-Lmut)
miR-Lmut

**Figure S1. The dosage response curves of the miR-L and miR-L$^{mut}$ -based targets and their dependencies of the TNRC6.** Related to Figure 1 and Figure 2.
**(A-B)** The performance of the miR-L-mediated 1-4×18nt constructs **(A)** and 1-4×19nt constructs **(B)**, measured by flow cytometry.
**(C-D)** The performance of the bulged target constructs either in the single copy **(C)** or in the multiple copy **(D)** configurations. The bulge designs were shown as the insets in the center of the plots. The bulge region starts from the 9th nucleotide on the target, and the lengths of which range from 1 nt to 5 nt.
**(E-F)** We performed flow cytometry on the cells transfected with the miR-L 4×17nt construct **(E)** or miR-L 1×21nt construct **(F)** described in **Figure 2B-C** along with the fluorescent protein-only negative control, the T6B peptide, or the catalytically dead T6B peptide (denoted as T6B$^{Mut}$), respectively. The design of the catalytically dead T6B peptide was consistent with the previous literature [S1].
**(G-J)** We performed flow cytometry on the cells co-transfected with the labeled constructs and the T6B peptide or the control described in **Figure 2B-C** to identify the TNRC6 dependence of different constructs. T6B restored the multiple-site targets to the similar level of the corresponding single-site targets.
**(K-N)** We performed flow cytometry on the cells transfected with the constructs regulated by the miR-L$^{mut}$, and compared them with the corresponding constructs regulated by the original miR-L. The inclusion of the central mismatch in the miRNA slightly enhanced the regulation.
**(O-P)** We performed flow cytometry to measure the TNRC6 dependence of the 1×21nt **(O)** and the 4×17nt **(P)** constructs, respectively. Similar TNRC6 dependence was observed for these miR-L$^{mut}$-regulated constructs compared to that of the original miR-L-regulated constructs shown in **Figure 2B-C**.

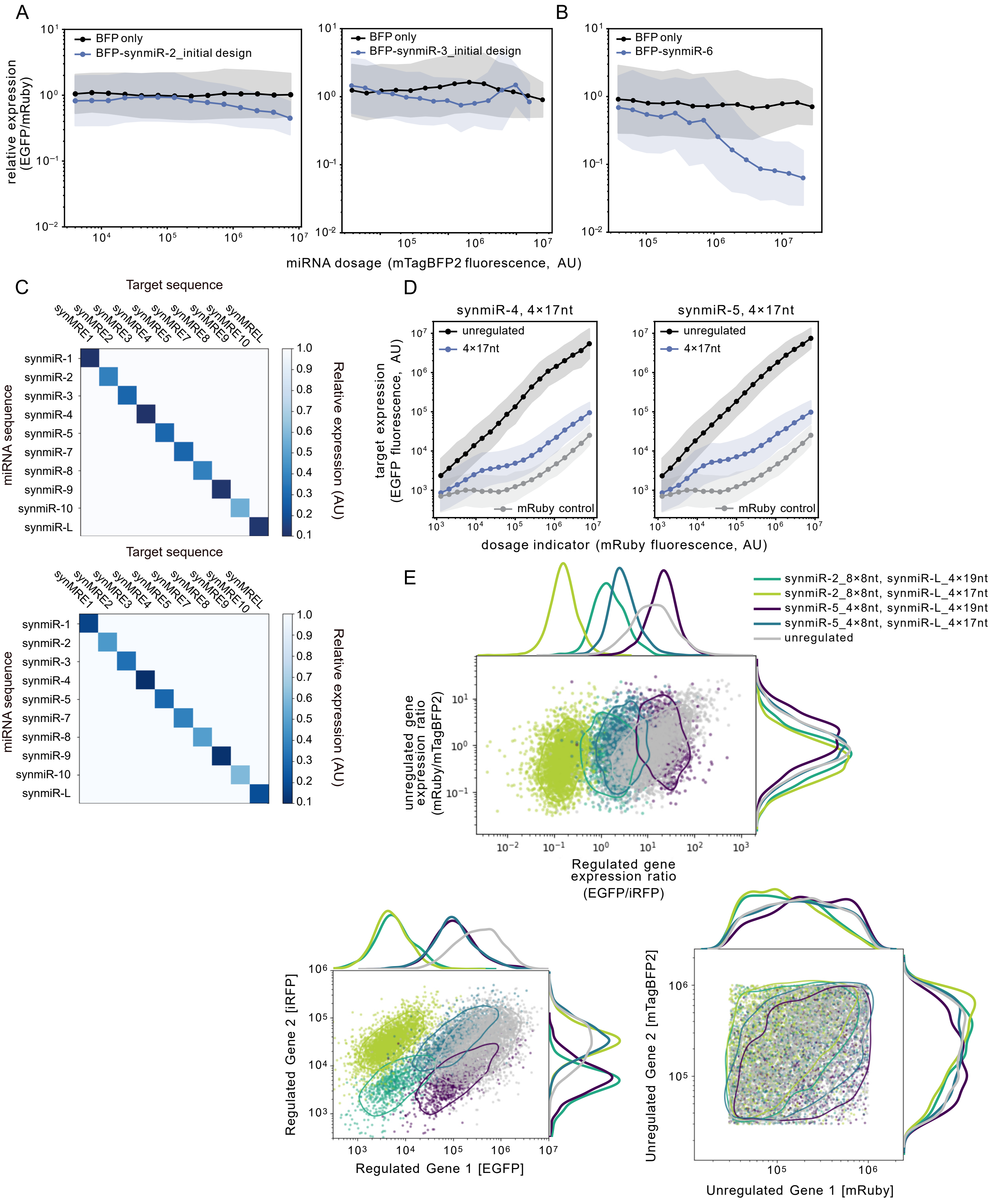

A
B
C
D
E
BFP only
BFP-synmiR-2_initial design
BFP-synmiR-3_initial design
BFP-synmiR-6
relative expression (EGFP/mRuby)
miRNA dosage (mTagBFP2 fluorescence, AU)
Target sequence
synMRE1 synMRE2 synMRE3 synMRE4 synMRE5 synMRE7 synMRE8 synMRE9 synMRE10 synMREL
synmiR-1
synmiR-2
synmiR-3
synmiR-4
synmiR-5
synmiR-7
synmiR-8
synmiR-9
synmiR-10
synmiR-L
miRNA sequence
Relative expression (AU)
synmiR-4, 4×17nt
synmiR-5, 4×17nt
unregulated
4×17nt
mRuby control
target expression (EGFP fluorescence, AU)
dosage indicator (mRuby fluorescence, AU)
synmiR-2_8×8nt, synmiR-L_4×19nt
synmiR-2_8×8nt, synmiR-L_4×17nt
synmiR-5_4×8nt, synmiR-L_4×19nt
synmiR-5_4×8nt, synmiR-L_4×17nt
unregulated
unregulated gene expression ratio (mRuby/mTagBFP2)
Regulated gene expression ratio (EGFP/iRFP)
Regulated Gene 2 [iRFP]
Regulated Gene 1 [EGFP]
Unregulated Gene 2 [mTagBFP2]
Unregulated Gene 1 [mRuby]

**Figure S2. The initial synmiR-2, 3, and 6 performances, the biological replicates of the orthogonality test, the 4×17nt designs of synmiR-4 and synmiR-5, and the measurements of the fluorescent protein expression of the dual-reporter system.** Related to Figure 4.
**(A-B)** We performed flow cytometry on the cells co-transfected with the circuit shown in **Figure 4A** using the initial sequence of synmiR-2 (**A, left panel**), synmiR-3 (**A, right panel**), synmiR-6 (**B**), and their corresponding targets with a single, fully complementary target site, respectively. The BFP only control does not have the 3'UTR miRNA. Relative expression levels were quantified as described in **Figure 4B**. Initial designs of synmiR-2 and synmiR-3 didn't function. synmiR-6 functioned but showed a sequence similarity to the endogenous miRNA hsa-mir-5697 (**STAR Methods**).
**(C)** The other two biological replicates of the experiment described by **Figure 4E**.
**(D)** The dosage response curves of 4×17nt designs of synmiR-4 and synmiR-5, measured by flow cytometry.
**(E)** The biological replicate of the experiment described in **Figure 4H**, along with the unregulated control group (grey). The unregulated group contains the dual reporter system, both without the regulation element. Other experimental settings are the same as described in **Figure 4H**.
Upper panel, the distributions of the ratio of both unregulated proteins ([mRuby]/[mTagBFP2]) against the ratio of both regulated proteins ([EGFP]/[iRFP]).
The bottom left panel shows the regulated proteins' distributions of each group. The bottom right panel shows the dosage indicators' distributions of each group.
The dosage indicators exhibit the same distribution among different groups, by contrast, different combinations of DIMMERs allow a clear separation of the populations by the regulated proteins. Additionally, DIMMERs tightly control the stoichiometry of the regulated proteins.

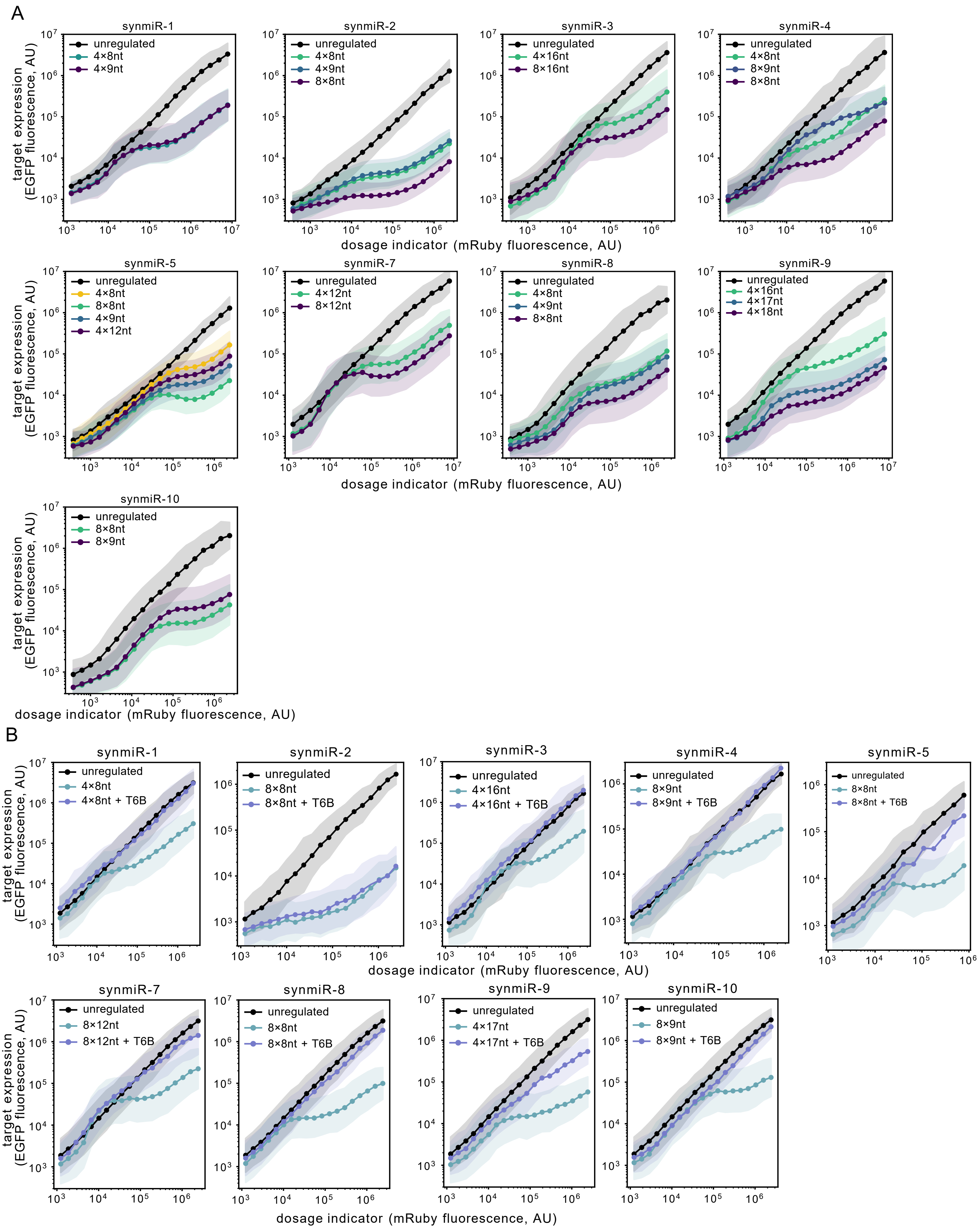
A
synmiR-1
unregulated
4×8nt
4×9nt
synmiR-2
unregulated
4×8nt
4×9nt
8×8nt
synmiR-3
unregulated
4×16nt
8×16nt
synmiR-4
unregulated
4×8nt
8×9nt
8×8nt
synmiR-5
unregulated
4×8nt
8×8nt
4×9nt
4×12nt
synmiR-7
unregulated
4×12nt
8×12nt
synmiR-8
unregulated
4×8nt
4×9nt
8×8nt
synmiR-9
unregulated
4×16nt
4×17nt
4×18nt
synmiR-10
unregulated
8×8nt
8×9nt
target expression (EGFP fluorescence, AU)
dosage indicator (mRuby fluorescence, AU)
B
synmiR-1
unregulated
4×8nt
4×8nt + T6B
synmiR-2
unregulated
8×8nt
8×8nt + T6B
synmiR-3
unregulated
4×16nt
4×16nt + T6B
synmiR-4
unregulated
8×9nt
8×9nt + T6B
synmiR-5
unregulated
8×8nt
8×8nt + T6B
synmiR-7
unregulated
8×12nt
8×12nt + T6B
synmiR-8
unregulated
8×8nt
8×8nt + T6B
synmiR-9
unregulated
4×17nt
4×17nt + T6B
synmiR-10
unregulated
8×9nt
8×9nt + T6B
target expression (EGFP fluorescence, AU)
dosage indicator (mRuby fluorescence, AU)

**Figure S3. A gallery of all the DIMMER designs based on different miRNAs and targets (A) and the dependence of TNRC6-based regulation (B).** Related to Figure 4.

Experimental settings are the same as described in **Figure 2B**. Almost all circuits used here relied on the TNRC6 to implement the inhibition, except synmiR-2 8×8nt, which might already be strong enough to achieve strong regulation.

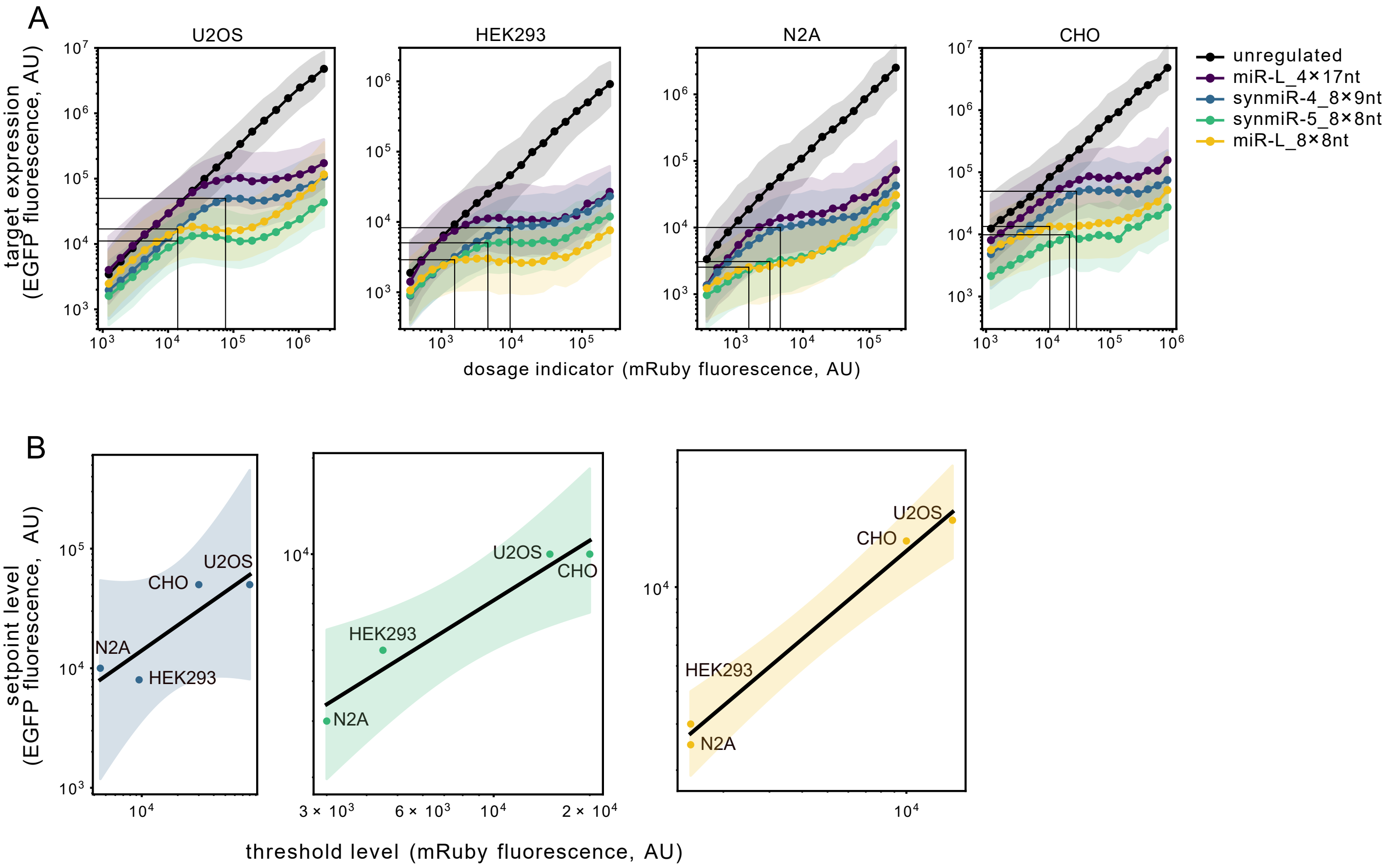


**Figure S4. IFFL works across different cell lines.** Related to Figure 5.
**(A)** The performance of different DIMMERs among various cell lines. The starting dosage of the dosage compensation behavior and the setpoint expression were indicated by the gray vertical lines and the horizontal lines, respectively.
**(B)** The threshold and the setpoint level co-vary across cell lines. The black solid line indicates a linear fit in the logarithm space (**STAR Methods**). See also **Figure 5B**.

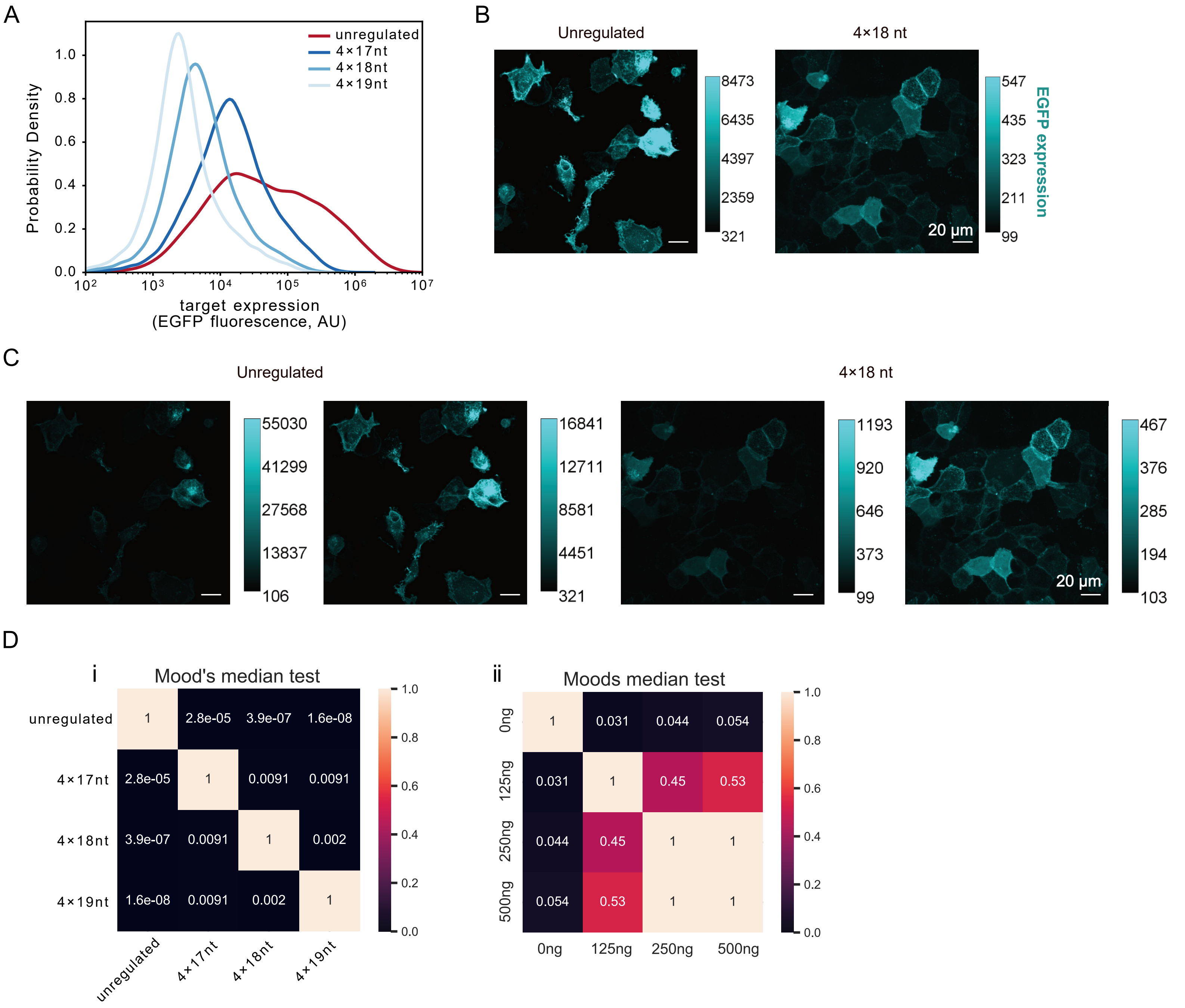


**Figure S5. DIMMER circuit improves the DNA-PAINT experiment.** Related to Figure 6.
**(A)** We performed flow cytometry on cells that were transfected with the EGFR-mEGFP with or without the DIMMER module. Cells were gated and binned by mRuby3 intensities. The expression of EGFR-mEGFP was plotted.
**(B-C)** Representative confocal microscopy images of the U2OS cells transfected with or without the DIMMER circuits **(B)** with different contrasts **(C,** indicated by the side color bars**)**. The images were taken using the 60× magnification objective. Numbers on the color bars indicate the fluorescence intensities measured by imageJ. Scale bar, 20 µm. DIMMERs allow a more uniform expression at a lower setpoint.
**(D)** The statistical test of the DNA-PAINT experiment described in **Figure 6E** (i) and **Figure 6F** (ii). The pipeline of the statistical test is described in the **STAR Methods** section.

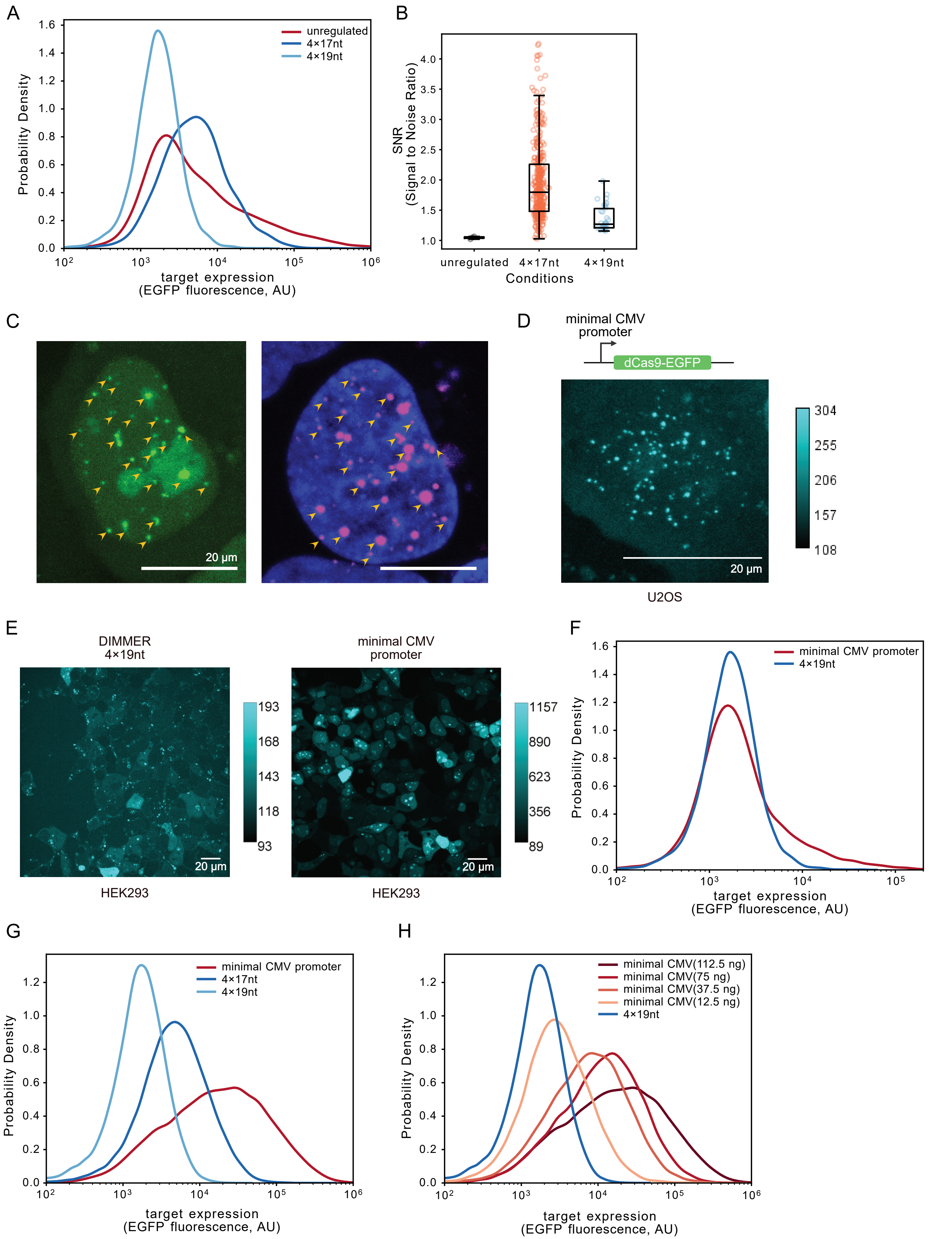

A
unregulated
4×17nt
4×19nt
Probability Density
target expression
(EGFP fluorescence, AU)
B
SNR
(Signal to Noise Ratio)
unregulated
4×17nt
4×19nt
Conditions
C
20 μm
D
minimal CMV
promoter
dCas9-EGFP
20 μm
U2OS
E
DIMMER
4×19nt
minimal CMV
promoter
20 μm
HEK293
F
minimal CMV promoter
4×19nt
G
minimal CMV promoter
4×17nt
4×19nt
H
minimal CMV(112.5 ng)
minimal CMV(75 ng)
minimal CMV(37.5 ng)
minimal CMV(12.5 ng)
4×19nt

**Figure S6. DIMMER circuit improves the CRISPR-based imaging.** Related to Figure 6.
**(A)** We performed flow cytometry on the cells co-transfected with the dCas9-EGFP and the guide RNA plasmid, with or without the DIMMER circuit regulating the dCas9-EGFP. The plot shows the distributions of the EGFP fluorescence signal.
**(B)** The boxplot shows the quantification of the signal-to-noise ratio (SNR) of the dots in the cells transfected with the dCas9-EGFP with or without the DIMMER module. Each scatter represents one dot inside a cell.
**(C)** The confocal microscopy images of one live U2OS cell expressing the dCas9-EGFP regulated by the 4×17nt circuit (left) and the same cell labeled by the DNA-FISH (right) probes targeting the telomeres post-fixation. The yellow arrowheads indicate the locations of the corresponding telomeres in the live cell and the fixed cell. Scale bar, 20 μm. Images were taken using the 60× objective. Detailed experimental procedures are described in **STAR Methods**.
**(D)** The confocal microscopy image of one live U2OS cell expressing the dCas9-EGFP driven by the minimal CMV promoter. Numbers on the color bars indicate the fluorescence intensities measured by imageJ. Scale bar, 20 μm.
**(E)** The confocal microscopy images of live HEK293T cells expressing the dCas9-EGFP either driven by the constitutive promoter and regulated by the 4×19nt DIMMER circuit (left) or driven by the minimal CMV promoter (right). Numbers on the color bars indicate the fluorescence intensities measured by imageJ. Scale bar, 20 μm.
**(F)** We performed flow cytometry on the U2OS cells expressing the dCas9-EGFP either driven by the constitutive promoter and regulated by the 4×19nt DIMMER circuit or driven by the minimal CMV promoter. The plot shows the distributions of the EGFP fluorescence signal.
**(G)** We performed flow cytometry on the HEK293T cells expressing the dCas9-EGFP either driven by the constitutive promoter and regulated by the 4×17nt and 4×19nt DIMMER circuits or driven by the minimal CMV promoter. The minimal CMV promoter generates a broader EGFP distribution.
**(H)** We performed flow cytometry on the HEK293T cells expressing the dCas9-EGFP either driven by the constitutive promoter and regulated by the 4×19nt DIMMER circuit or driven by the minimal CMV promoter with different transfection doses. The lowest transfection amount condition (12.5 ng) still cannot generate a distribution matching that of the 4×19nt DIMMER circuit.

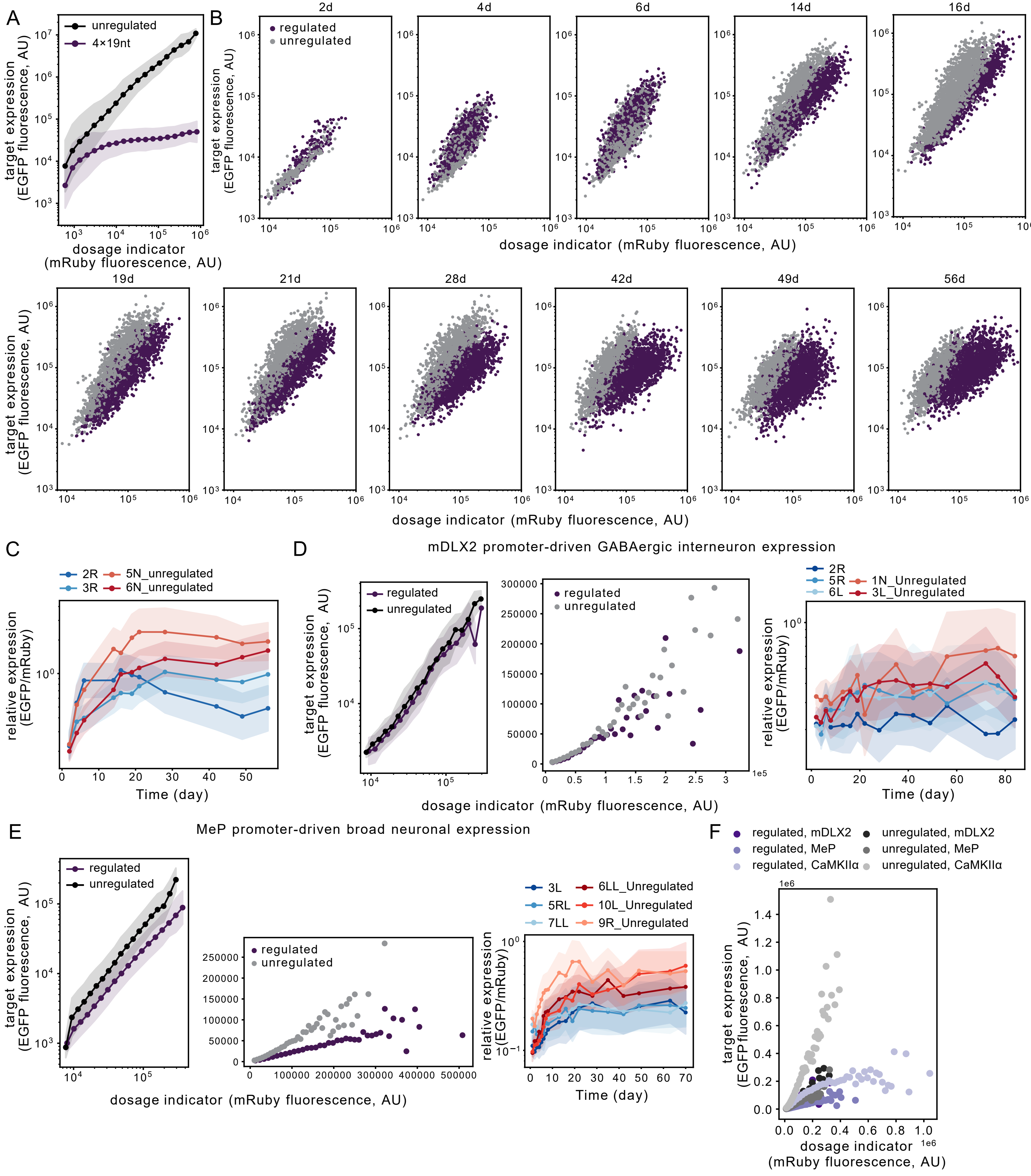
A
unregulated
4×19nt
target expression (EGFP fluorescence, AU)
dosage indicator (mRuby fluorescence, AU)
B
2d
4d
6d
14d
16d
regulated
unregulated
19d
21d
28d
42d
49d
56d
target expression (EGFP fluorescence, AU)
dosage indicator (mRuby fluorescence, AU)
C
2R
3R
5N_unregulated
6N_unregulated
relative expression (EGFP/mRuby)
Time (day)
D
mDLX2 promoter-driven GABAergic interneuron expression
regulated
unregulated
target expression (EGFP fluorescence, AU)
dosage indicator (mRuby fluorescence, AU)
2R
5R
6L
1N_Unregulated
3L_Unregulated
relative expression (EGFP/mRuby)
Time (day)
E
MeP promoter-driven broad neuronal expression
regulated
unregulated
target expression (EGFP fluorescence, AU)
dosage indicator (mRuby fluorescence, AU)
3L
5RL
7LL
6LL_Unregulated
10L_Unregulated
9R_Unregulated
relative expression (EGFP/mRuby)
Time (day)
F
regulated, mDLX2
regulated, MeP
regulated, CaMKIIα
unregulated, mDLX2
unregulated, MeP
unregulated, CaMKIIα
target expression (EGFP fluorescence, AU)
dosage indicator (mRuby fluorescence, AU)

**Figure S7. DIMMER circuits operate in live mouse brains.** Related to Figure 7.
**(A)** We performed flow cytometry on the U2OS cells that were co-transfected with the circuits described in **Figure 7G**, upper panel.
**(B)** We quantified the expressions of H2B-mRuby3 and H2B-EGFP driven by the CaMKIIα promoter which labels the excitatory neurons over time. Each dot represents the protein expressions of one single cell. Expressions of the unregulated and the regulated groups both accumulated with time, and gradually showed separation from each other. The time shown on the top of each plot indicates the time post injection. The image analysis procedure is described in detail in **STAR Methods**.
**(C)** We plotted the relative expression (the median single-cell EGFP/mRuby3 value) of each mouse at each single time point in the CaMKIIα promoter cohort. The orange curve and the red curve show the EGFP/mRuby dynamics of two mice in the unregulated group, and the light blue and the dark blue curves show the EGFP/mRuby dynamics of two mice in the 4×19nt DIMMER-regulated group. The shaded region was calculated as described in **Figure 1F**.
**(D)(E)** We used the mDLX2 promoter to drive the GABAergic interneuronal expression **(D)**, and used the MeP promoter to drive pan-neuronal expression **(E)** in the mouse brains. The left and the middle panels show the quantification of the pooled cells under each experimental condition, as described in **Figure 7H**, with the corresponding tissue-specific promoter, respectively. The right panels show the relative expression of each mouse at each single time point, as described in **(C)**, with the corresponding tissue-specific promoter, respectively.
**(F)** We pooled all the data of different cohorts together and plotted the EGFP intensities against mRuby3 intensities in the linear scale. The dots were calculated by the binned mRuby3 intensities.

**Data S1 Mathematical modeling of the miRNA-based gene dosage compensation circuits, related to Figure 1.**

### Supplemental Modeling Text

#### 1 Modeling miRNA-based gene dosage compensation circuits

Here we develop a simplified mathematical model of miRNA-based incoherent feed-forward loop (IFFL) circuits, and use it to explore how various parameters control the scaling of target gene expression with gene dosage. The focus in this section is to understand how general properties of the circuit influence its ability to perform dosage compensation. We also include a second Supplementary modeling section that focuses on the role of specific molecular mechanisms.

In this model, we focused on three sets of dynamic processes: RISC production and removal, mRNA production and removal, and RISC-mRNA complex formation.

**1. RISC production and removal.** We assume miRNA is expressed, processed, and loaded into Argonaute (Ago) proteins to form an active (mature) RNA-Induced Silencing Complex (RISC) [S2]. We denote the concentration of mature RISC (containing the miRNA) as $r$. We assume RISC is produced at a total rate of $D\beta_{\mathrm{r}}$, where $D$ denotes gene copy number (gene dosage) and $\beta_{\mathrm{r}}$ denotes the rate of production RISC production per gene copy. This expression implicitly assumes that miRNA expression levels do not saturate available miRNA processing machinery, Ago, or other components. We also assume that the RISC complex is removed at total rate $\gamma_{\mathrm{r}} r$, where $\gamma_{\mathrm{r}}$ denotes a combined rate constant for dilution, degradation, and other removal processes.

**2. mRNA production and removal.** We assume mRNA, denoted $m$, is produced at a rate proportional to gene copy number, $D\beta_{\mathrm{m}}$, where $\beta_{\mathrm{m}}$ is the mRNA production rate per gene copy. mRNA can also be removed at a rate $\gamma_{\mathrm{m}} m$ due to dilution and degradation. We note that even though mRNA and miRNA are produced from the same engineered locus, the production rate constants $\beta_{\mathrm{m}}$ and $\beta_{\mathrm{r}}$ can differ, since the miRNA and mRNA are produced from distinct promoters, and/or are processed through different downstream pathways.

**3. RISC-mRNA complex formation and dissociation.** We assume that the RISC and target mRNA associate to form a complex, whose concentration is denoted $C$, at a rate $k_{\mathrm{on}} rm$, following mass action kinetics with rate constant, $k_{\mathrm{on}}$. Once formed, this complex can dissociate at rate $k_{\mathrm{off}} C$, undergo catalytic mRNA degradation, at rate $k_{\mathrm{c}} C$.

These chemical reactions can be summarized as follows, using $G$ to denote the gene, present at copy number (dosage) $D$:

$$
\begin{aligned}
G &\xrightarrow{D\beta_{\mathrm{r}}} G + r \\
r &\xrightarrow{\gamma_{\mathrm{r}}} \emptyset \\
G &\xrightarrow{D\beta_{\mathrm{m}}} G + m \\
m &\xrightarrow{\gamma_{\mathrm{m}}} \emptyset \\
r + m &\xrightarrow{k_{\mathrm{on}}} C \\
C &\xrightarrow{k_{\mathrm{off}}} r + m \\
C &\xrightarrow{k_{\mathrm{c}}} r
\end{aligned}
$$

With these definitions and assumptions, we can write down a set of ordinary differential equations for the three variables, $r$, $m$, and $C$.

$$
\begin{aligned}
\frac{dr}{dt} &= \beta_{\mathrm{r}} D - \gamma_{\mathrm{r}} r - k_{\mathrm{on}} rm + (k_{\mathrm{c}} + k_{\mathrm{off}})C \\
\frac{dm}{dt} &= \beta_{\mathrm{m}} D - \gamma_{\mathrm{m}} m - k_{\mathrm{on}} rm + k_{\mathrm{off}} C \\
\frac{dC}{dt} &= k_{\mathrm{on}} rm - (k_{\mathrm{c}} + k_{\mathrm{off}})C
\end{aligned}
$$

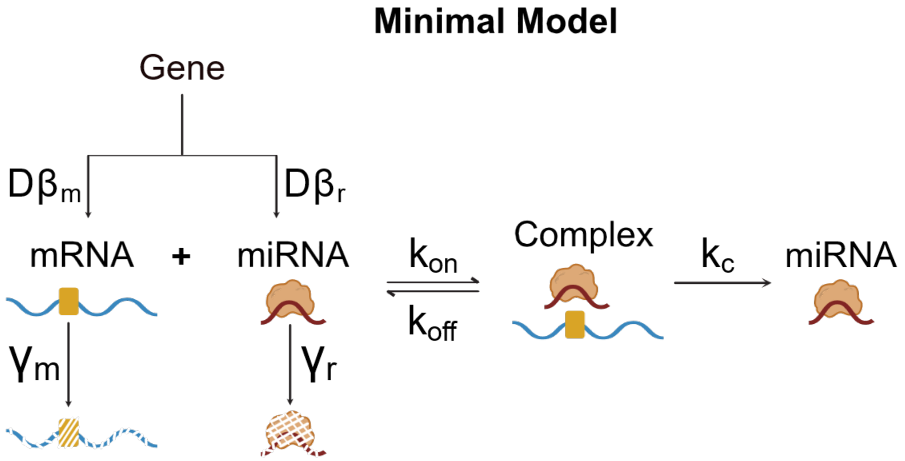


**Data S1-Figure 1: Reactions in the minimal model,** including mRNA and miRNA production and removal, as well as RISC complex formation and catalytic mRNA degradation. Parameters on arrows denote reaction rates.

These equations resemble previous modeling of regulation by small RNAs [S3, S4], except for the explicit incorporation of gene dosage in the control of both mRNA and miRNA.

To gain insight into the possible behaviors of this system, we first non-dimensionalize it. We define a dimensionless time, $\widetilde{t} = \gamma_{\mathrm{r}} t$, by rescaling time in units of the RISC lifetime. We also define a dimensionless RISC concentration, $\widetilde{r} = r/(\beta_{\mathrm{r}}/\gamma_{\mathrm{r}})$. This effectively rescales $r$ in units of the unregulated steady-state expression level produced by a single copy of the gene. We similarly define a dimensionless mRNA concentration, $\widetilde{m} = m/(\beta_{\mathrm{m}}/\gamma_{\mathrm{m}})$, normalizing $m$ by its single copy steady state expression level. Finally, we define a dimensionless concentration of the RISC-miRNA-mRNA complex, $\widetilde{C} = C/(\beta_{\mathrm{m}}/\gamma_{\mathrm{m}})$.

In addition, we define a set of convenient dimensionless parameter ratios:

$$\gamma = \frac{\gamma_{\mathrm{m}}}{\gamma_{\mathrm{r}}}$$
$$\beta = \frac{\beta_{\mathrm{m}}}{\beta_{\mathrm{r}}}$$
$$\widetilde{k}_{\mathrm{on}} = \frac{k_{\mathrm{on}}\beta_{\mathrm{r}}}{\gamma_{\mathrm{m}}\gamma_{\mathrm{r}}}$$
$$\widetilde{k}_{\mathrm{off}} = \frac{k_{\mathrm{off}}}{\gamma_{\mathrm{m}}}$$
$$\widetilde{k}_{\mathrm{c}} = \frac{k_{\mathrm{c}}}{\gamma_{\mathrm{m}}}$$
$$K = \frac{\widetilde{k}_{\mathrm{c}} + \widetilde{k}_{\mathrm{off}}}{\widetilde{k}_{\mathrm{on}}}$$

In the non-dimensionalized system, the differential equations can be written as,

$$\frac{d\widetilde{r}}{d\widetilde{t}} = D - \widetilde{r} - \widetilde{k}_{\mathrm{on}}\beta\widetilde{m}\widetilde{r} + (\widetilde{k}_{\mathrm{c}} + \widetilde{k}_{\mathrm{off}})\beta\widetilde{C}$$
$$\gamma^{-1}\frac{d\widetilde{m}}{d\widetilde{t}} = D - \widetilde{m} - \widetilde{k}_{\mathrm{on}}\widetilde{m}\widetilde{r} + \widetilde{k}_{\mathrm{off}}\widetilde{C}$$
$$\gamma^{-1}\frac{d\widetilde{C}}{d\widetilde{t}} = \widetilde{k}_{\mathrm{on}}(\widetilde{m}\widetilde{r} - K\widetilde{C})$$

By definition, at steady state, the time derivatives all equal zero. Denoting steady values with a subscript $s$, we then have:

$$D - \widetilde{r}_{\mathrm{s}} - \widetilde{k}_{\mathrm{on}}\beta\widetilde{m}_{\mathrm{s}}\widetilde{r}_{\mathrm{s}} + (\widetilde{k}_{\mathrm{c}} + \widetilde{k}_{\mathrm{off}})\beta\widetilde{C}_{\mathrm{s}} = 0$$
$$D - \widetilde{m}_{\mathrm{s}} - \widetilde{k}_{\mathrm{on}}\widetilde{m}_{\mathrm{s}}\widetilde{r}_{\mathrm{s}} + \widetilde{k}_{\mathrm{off}}\widetilde{C}_{\mathrm{s}} = 0$$
$$\widetilde{m}_{\mathrm{s}}\widetilde{r}_{\mathrm{s}} - K\widetilde{C}_{\mathrm{s}} = 0$$

Solving the equations above, we obtain an equation for steady-state mRNA concentration:

$$\widetilde{m}_{\mathrm{s}} = \frac{D}{1 + \frac{\widetilde{k}_{\mathrm{c}}}{K}D}$$

Henceforth, we will omit the tildes for notational convenience, and switch to the non-dimensionalized variables and parameters.

In the limit of large dosage, $\frac{k_c}{K}D \gg 1$. As a result, $m_s$ approaches a limiting value, $m_s \to \frac{K}{k_c}$ independent of gene dosage. This is the regime that the constructs developed here are targeting.

Main text Figure 1D plots this expression, and Data S1-Figure 2A and B plot the expression in different parameters, i.e. $k_{\text{off}}$, and $k_c$.

Finally, to estimate biological parameter values, we assume that the miRNA and mRNA are produced at similar rates, using estimated parameter values derived from [S5] with tunable ranges, as listed in the following table:

| **Non-dimensional parameter** | **Dimensionless values** |
|---|---|
| $\beta$ | 1 |
| $\gamma$ | 0.8 |
| $k_{\text{on}}$ | 200000 |
| $k_{\text{off}}$ | 10, 100, 1000, 10000 |
| $k_c$ | 400, 40, 4, 0.4, 0.004, 0.0004 |

This simple model suggests that steady state mRNA concentration can—under some regimes—achieve dosage independence.

#### 1.1 Incorporating ultrasensitivity

Multiple mechanisms can give rise to ultrasensitivity in miRNA regulation, which is not present in the model so far. To allow for ultrasensitivity, we considered a more general, phenomenological model in which target mRNA inhibition by miRNA follows a Hill function of the RISC concentration, with Hill coefficient $n$. This treatment omits intermediate steps, i.e., the RISC-mRNA complex formation and dissociation. With this assumption, we can write down a different set of ordinary differential equations for the two variables, $r$, and $m$.

$$\frac{dr}{dt} = \beta_r D - \gamma_r r$$
$$\frac{dm}{dt} = \frac{\beta_m D}{1 + (\frac{r}{\kappa})^n} - \gamma_m m$$

To non-dimensionalize this system, we define $\widetilde{t} = \gamma_r t$, $\widetilde{r} = r/(\beta_r/\gamma_r)$, and $\widetilde{m} = m/(\beta_m/\gamma_m)$. Additionally, we define $\widetilde{K} = \frac{\beta_r}{\gamma_r \kappa}$ for convenience. The steady state expression of mRNA concentration can then be written as:

$$\widetilde{m_s} = \frac{D}{1 + (\widetilde{K}D)^n}$$

Data S1-Figure 2C plots this expression when $\widetilde{K} = 1$ and $n = 0.5, 1, 2$. Critically, only when $n = 1$, does $\widetilde{m}_s$ approach a limiting value independent of $D$. If $n < 1$, $\widetilde{m}_s$ shows a sublinear increase with $D$. If $n > 1$, $\widetilde{m}_s$ shows a biphasic dependence on $D$ (Data S1-Figure 2C).

This analysis suggests that it is important to maintain a Hill coefficient of $n \sim 1$ for optimal dosage compensation.

#### 1.2 Incorporating complex degradation and the bounded mRNA

Thus far, we ignored possible degradation of mRNA and miRNA within the complex, and did not explicitly account for total mRNA, denoted $m_{\text{tot}}$, which includes both free mRNA and mRNA engaged in RISCs. Here, we incorporate these additional reactions and analyze their effects on the dosage response behavior. Specifically, we added two additional reactions:

$$C \xrightarrow{\gamma_r} m$$
$$C \xrightarrow{\gamma_m} r$$

We now write down the modified ordinary differential equations:

$$\frac{dr}{dt} = \beta_r D - \gamma_r r - k_{\text{on}} rm + (k_c + k_{\text{off}} + \gamma_m)C$$
$$\frac{dm}{dt} = \beta_m D - \gamma_m m - k_{\text{on}} rm + (k_{\text{off}} + \gamma_r)C$$
$$\frac{dC}{dt} = k_{\text{on}} rm - (k_c + k_{\text{off}} + \gamma_m + \gamma_r)C$$

We then perform non-dimensionalization, as above. The only dimensionless parameter that has been changed is $K$, which now is $K = \frac{\widetilde{k}_c + \widetilde{k}_{\text{off}} + 1 + \gamma^{-1}}{\widetilde{k}_{\text{on}}}$, leading to the following dimensionless differential equations:

$$\frac{d\widetilde{r}}{d\widetilde{t}} = D - \widetilde{r} - \widetilde{k}_{\text{on}} \beta \widetilde{m}\widetilde{r} + (\widetilde{k}_c + \widetilde{k}_{\text{off}} + 1)\beta \widetilde{C}$$
$$\gamma^{-1}\frac{d\widetilde{m}}{d\widetilde{t}} = D - \widetilde{m} - \widetilde{k}_{\text{on}} \widetilde{m}\widetilde{r} + (\widetilde{k}_{\text{off}} + \gamma^{-1})\widetilde{C}$$
$$\gamma^{-1}\frac{d\widetilde{C}}{d\widetilde{t}} = \widetilde{k}_{\text{on}}(\widetilde{m}\widetilde{r} - K\widetilde{C})$$

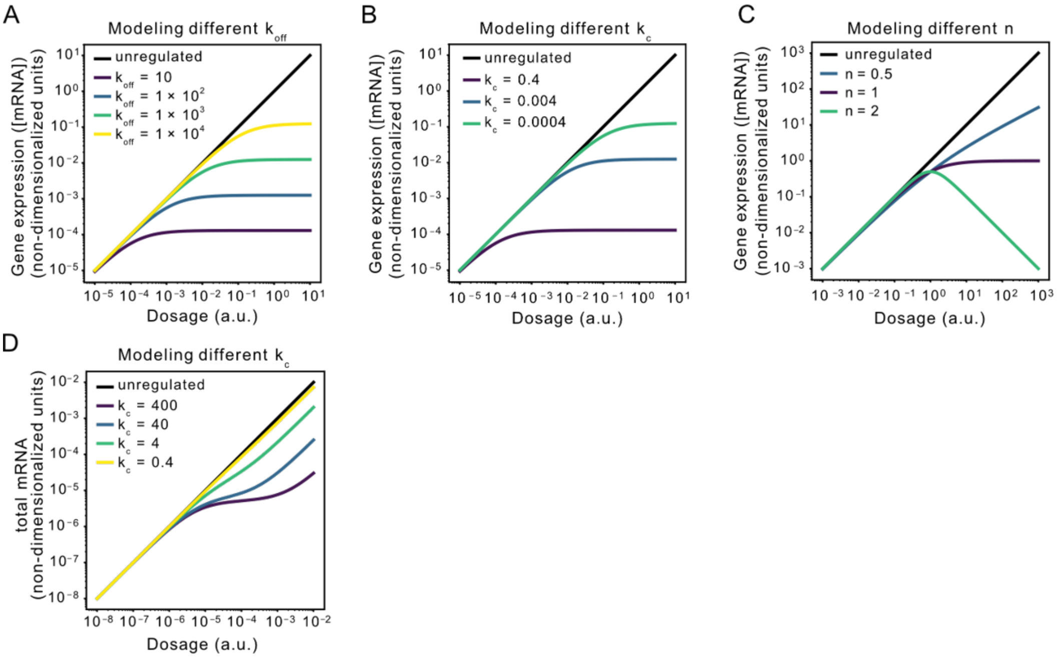


**Data S1-Figure 2: The minimal model of the miRNA-mediated IFFL.** A, modeling the steady-state, dimensionless mRNA concentration under different $k_{\mathrm{off}}$ while maintaining $k_{\mathrm{on}} = 2 \times 10^5, k_{\mathrm{c}} = 0.4$. B, modeling the steady-state, dimensionless mRNA concentration under different $k_{\mathrm{c}}$ while maintaining $k_{\mathrm{on}} = 2 \times 10^5, k_{\mathrm{off}} = 10$. C, modeling the steady-state, dimensionless mRNA concentration under different Hill coefficient n while maintaining $K = 1$. D, modeling the steady-state, dimensionless total mRNA amount (including the free mRNA and the bounded complex) under different $k_{\mathrm{c}}$ while maintaining $k_{\mathrm{on}} = 2 \times 10^5, k_{\mathrm{off}} = 10$ by considering the complex degradation caused by natural degradation of either mRNA or miRNA, and the translational contribution of the bounded complex.

From this, we can compute steady state concentrations:

$$\widetilde{C_{\mathrm{s}}} = \frac{\widetilde{m_{\mathrm{s}}}\widetilde{r_{\mathrm{s}}}}{K}$$

$$\widetilde{m_{\mathrm{s}}} = \frac{D}{1 + \frac{(1+\widetilde{k_{\mathrm{c}}})\widetilde{r_{\mathrm{s}}}}{K}}$$

$$\widetilde{r_{\mathrm{s}}} = \frac{D(\widetilde{k_{\mathrm{c}}}+1) - \beta D\gamma^{-1} - K + \sqrt{(D(\widetilde{k_{\mathrm{c}}}+1) - \beta D\gamma^{-1} - K)^2 + 4DK(\widetilde{k_{\mathrm{c}}}+1)}}{2(\widetilde{k_{\mathrm{c}}}+1)}$$

Considering the parameter regime where $\widetilde{k_{\mathrm{c}}} + 1 \gg \beta\gamma^{-1}$, the $\beta D\gamma^{-1}$ term is negligible. Therefore, we get $\widetilde{r_{\mathrm{s}}} \approx D$. Henceforth, we drop the tildes for notational convenience, as in the previous sections. We write down the expression for $m_{\mathrm{tot}}$:

$$m_{\mathrm{tot}} = m_{\mathrm{s}} + C_{\mathrm{s}} = m_{\mathrm{s}}(1 + \frac{r_{\mathrm{s}}}{K}) = \frac{D(K + r_{\mathrm{s}})}{K + (k_{\mathrm{c}}+1)r_{\mathrm{s}}} \approx \frac{D(K+D)}{K+(k_{\mathrm{c}}+1)D}$$

We now discuss the behavior of $m_{\mathrm{tot}}$ at different dosages:

1. When $D \ll \frac{K}{k_{\mathrm{c}}+1} < K$: $m_{\mathrm{tot}} \to D$. In this case, the total mRNA scales linearly with the dosage, similar to the small dosage regime in the minimal model.

2. When $\frac{K}{k_{\mathrm{c}}+1} < D \ll K$: $m_{\mathrm{tot}} \approx \frac{D}{1+\frac{(k_{\mathrm{c}}+1)D}{K}} \to \frac{K}{k_{\mathrm{c}}+1}$. In this case, $m_{\mathrm{tot}}$ shows dosage compensation.

3. When $D \gg K$: $m_{\mathrm{tot}} \to \frac{D}{k_{\mathrm{c}}+1}$. In this case, the total mRNA is again approximately proportional to dosage, but has a lower setpoint compared with the unregulated condition due to repression. Data S1-Figure 2D depicts this behavior.

### 2 Conclusions from simple modeling of miRNA-based gene dosage compensation circuits

Based on the results above, we obtain the following main design guidelines for miRNA-based dosage compensation circuits:

1. miRNA-based circuits incorporating miRNA-mRNA complex formation and catalytic mRNA degradation should be able to achieve dosage compensation.

2. However, a critical requirement for dosage compensation is non-ultrasensitive regulation of mRNA by miRNA, i.e. a Hill coefficient $n \sim 1$.

3. Degradation of mRNA and miRNA within the RISC can affect dosage compensation. However, these reactions still allow dosage compensation within a limited regime provided that $k_c$ is sufficiently high (Data S1-Figure 2D). The cutoff for

this behavior occurs when dosage levels increase enough to make $r_s$ becomes comparable to $K$, leading to a "tail" in the expression versus dosage plot.

**Data S2** Iterative engineering, auxiliary analyses of variance, alternative visualizations of the main figure datasets, and other technical controls, related to Figure 2, 3, 4, 5, 7.

Data S2-1

1st generation of circuits: co-transcribed miRNA design

① miRNA in the 3'UTR

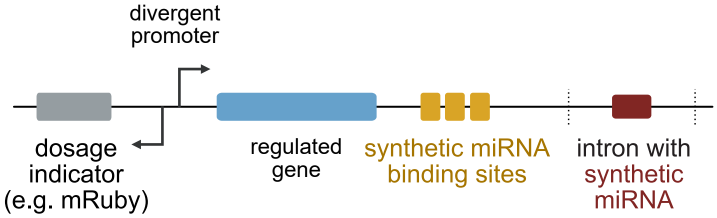


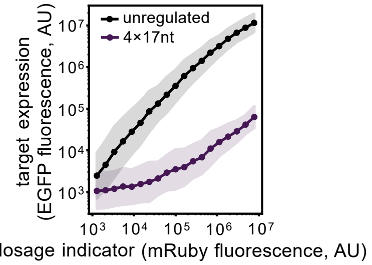


② miRNA in the 5'UTR

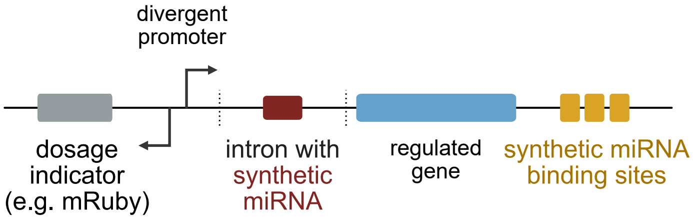


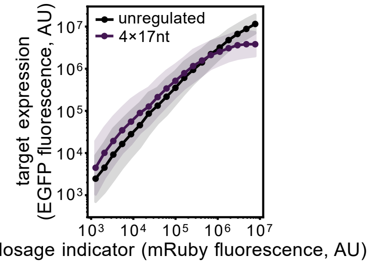


③ miRNA inside the transcript

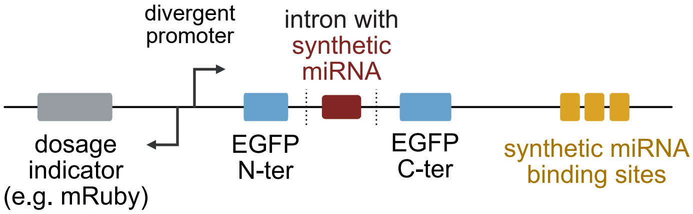


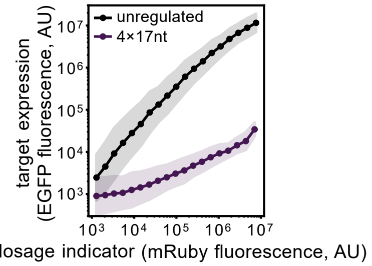


2nd generation of circuits: separately transcribed miRNA design

① miRNA in the 3'UTR

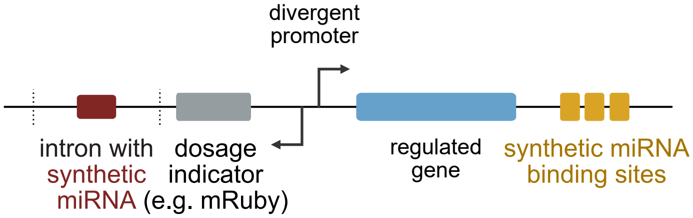


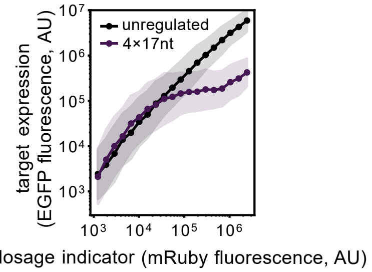


② miRNA in the 5'UTR

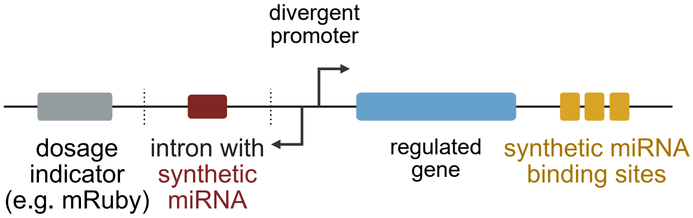


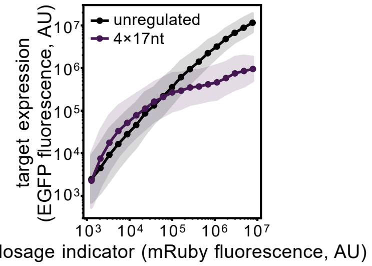


③ miRNA inside the transcript

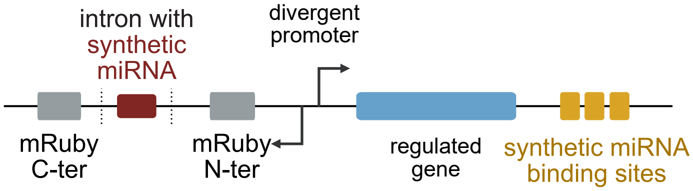


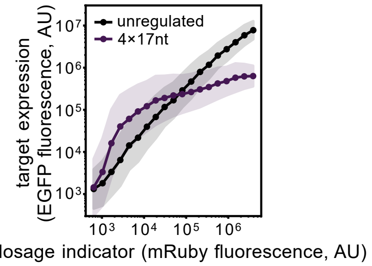


3rd generation of circuits: co-transcribed miRNA design, optimized splicing sequence

① miRNA in the 3'UTR

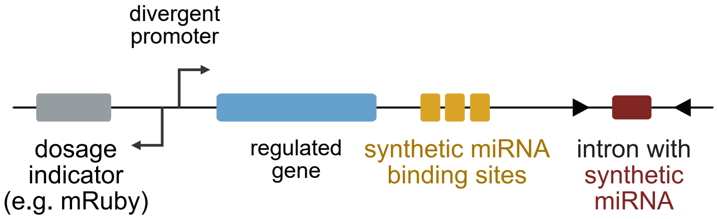


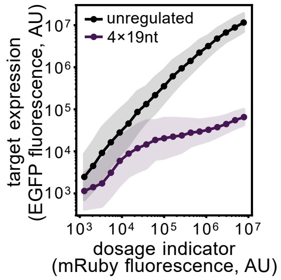


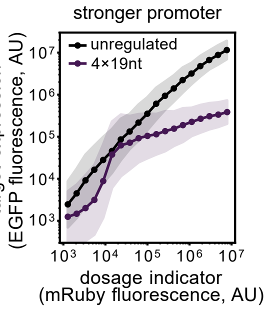

**Data S2-1. The iterative engineering process of different circuit architectures, related to Figure 2.** Left panel, various circuit configurations; right panel, corresponding circuit quantitative behaviors measured by flow cytometry.

In the first generation of the circuit, a co-transcribed miRNA design was used. Placing the intronic synthetic miRNA in the 3'-UTR or in the middle of the transcript both lead to the strong repression effects that might be caused by the destabilization of the transcript that is independent of the miRNA inhibition. Placing the intronic synthetic miRNA in the 5'-UTR of the transcript does not produce enough miRNA.

In the second generation of the circuit, the intronic synthetic miRNA sequence was placed in the separate transcript (which is mRuby3). No matter where the miRNA sequence is inserted, the dosage compensation behavior is maintained. However, when the miRNA sequence is placed into the 3'-UTR, the amount of the mature miRNA seems to be the greatest, since the circuit has the lowest setpoint.

In the third generation of the circuit, the intronic synthetic miRNA sequence was placed in the same transcript as the regulated EGFP. The key difference between the third generation design and the first generation design is that the splicing signal sequence is optimized (indicated by the black triangles). The dosage compensation behavior is achieved, and the utilization of a stronger promoter changes the setpoint.

**Data S2-1 Table: Summary of performances of different circuit configurations.**

| **Circuit configuration** | **miRNA production** | **Transcript stability** |
|---|---|---|
| 1st generation, ① | normal-to-low miRNA production | unstable |
| 1st generation, ② | low miRNA production | stable |
| 1st generation, ③ | almost no miRNA production | unstable |
| 2nd generation, ① | normal miRNA production | stable |
| 2nd generation, ② | lower miRNA production compared to 2nd generation, ① | stable |
| 2nd generation, ③ | lower miRNA production compared to 2nd generation, ① | stable |
| 3rd generation, ① | normal miRNA production | stable |
| 3rd generation, ①, stronger promoter | normal miRNA production | stable |

Data S2-2

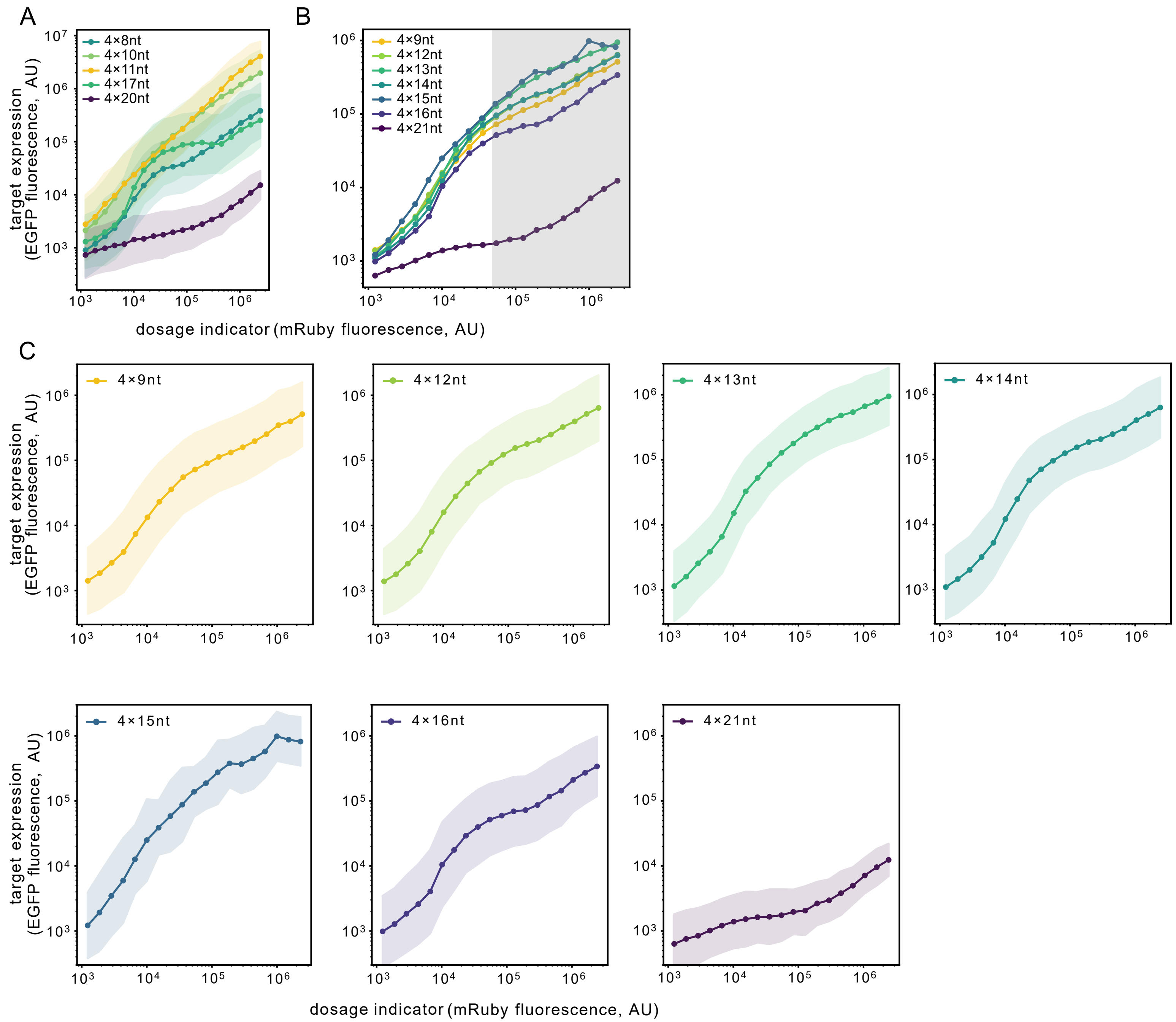


**Data S2-2. The dosage response curves of multiple miR-L 4×n circuits, related to Figure 3.**

**(A)** The curves in **Figure 3A** right panels with the geometric variances plotted.

**(B)** We performed flow cytometry on the cells transfected with the miR-L regulating 4×n circuits described in **Figure 3A**. Gray rectangle indicates the gated region of the **Figure 3A** left panel.

**(C)** The curves in **(B)** with the geometric variances were plotted separately to avoid being cluttered together.

Data S2-3

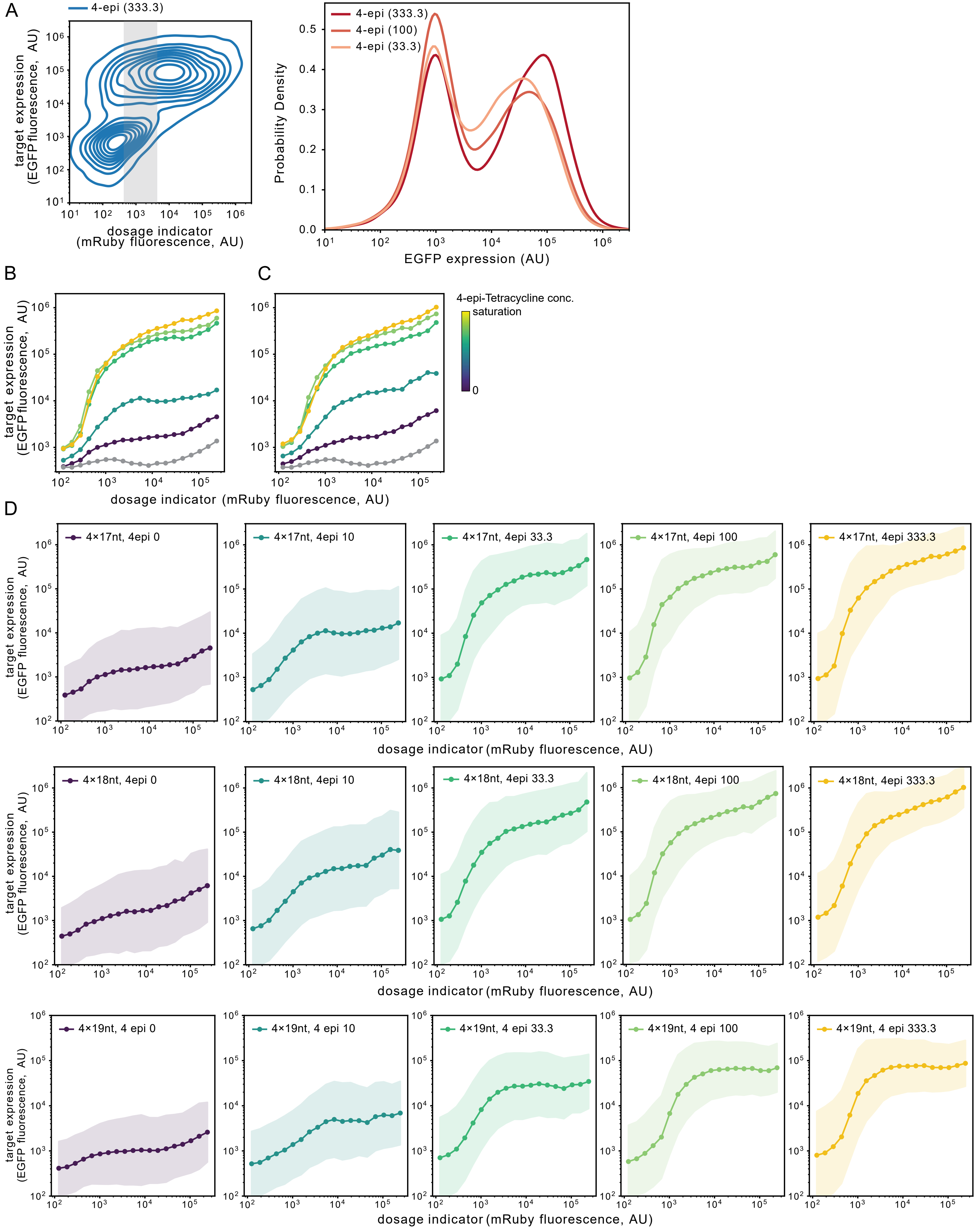
A
4-epi (333.3)
target expression (EGFP fluorescence, AU)
dosage indicator (mRuby fluorescence, AU)
4-epi (333.3)
4-epi (100)
4-epi (33.3)
Probability Density
EGFP expression (AU)
B
C
target expression (EGFP fluorescence, AU)
4-epi-Tetracycline conc.
saturation
0
dosage indicator (mRuby fluorescence, AU)
D
4×17nt, 4epi 0
4×17nt, 4epi 10
4×17nt, 4epi 33.3
4×17nt, 4epi 100
4×17nt, 4epi 333.3
4×18nt, 4epi 0
4×18nt, 4epi 10
4×18nt, 4epi 33.3
4×18nt, 4epi 100
4×18nt, 4epi 333.3
4×19nt, 4 epi 0
4×19nt, 4 epi 10
4×19nt, 4 epi 33.3
4×19nt, 4 epi 100
4×19nt, 4 epi 333.3
target expression (EGFP fluorescence, AU)
dosage indicator (mRuby fluorescence, AU)

**Data S2-3. The dosage response curves of the inducible DIMMER circuits in TRex cells, related to Figure 3.**

**(A)** The left panel shows the contour plot of the 4×19nt construct described in **Figure 3E**. The concentration of 4-epi was 333.3 ng/mL. Cells falling into the shaded grey region were selected to plot the EGFP distribution in the right panel. Cells in **Figure 3E** falling into the same mRuby3 range were also selected to plot the EGFP distribution at the 4-epi concentration of 100 and 33.3 ng/mL. The bimodal distribution of EGFP indicates the bursty nature of the Tet-On promoter used here, and explains the seemingly ultrasensitive behavior observed in **Figure 3E** when the dosage is low.

**(B-C)** We performed flow cytometry on the TRex cell line which was transfected with the 4×17nt **(B)** or 4×18nt **(C)** DIMMER construct described in **Figure 3E**. The concentrations of the 4-epi-Tetracycline, from purple to yellow, were 0, 10, 33.3, 100, 333.3 ng/mL. The gray curve denotes the mRuby-only transfection control.

**(D)** The curves in **Figure 3E** and **Data S2-3 B-C** with the geometric variances plotted separately to avoid being cluttered together.

Data S2-4

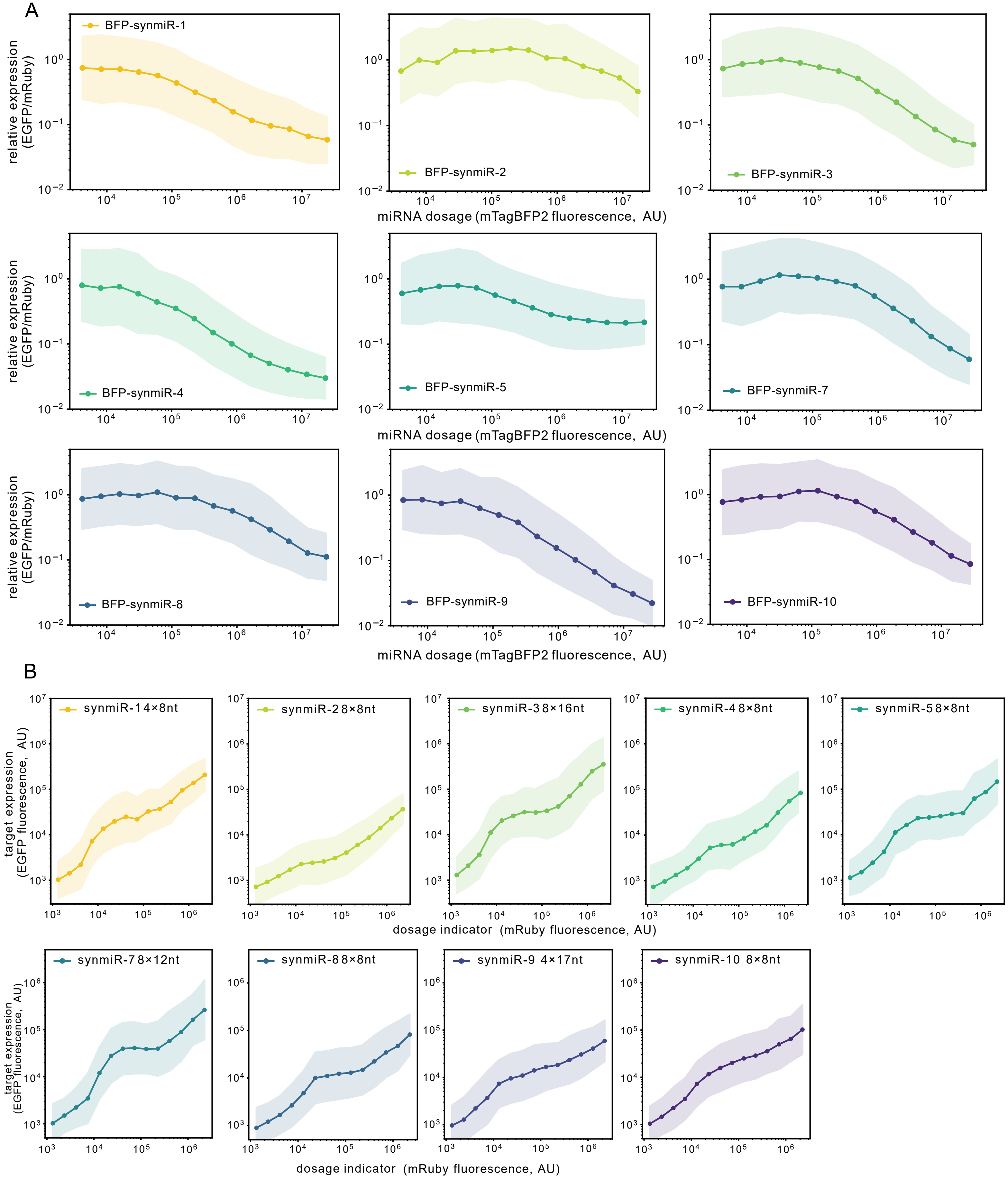


**Data S2-4. The geometric variance of the dosage response curves in Figure 4D (A) and Figure 4F (B) to avoid curve cluttering, related to Figure 4.**

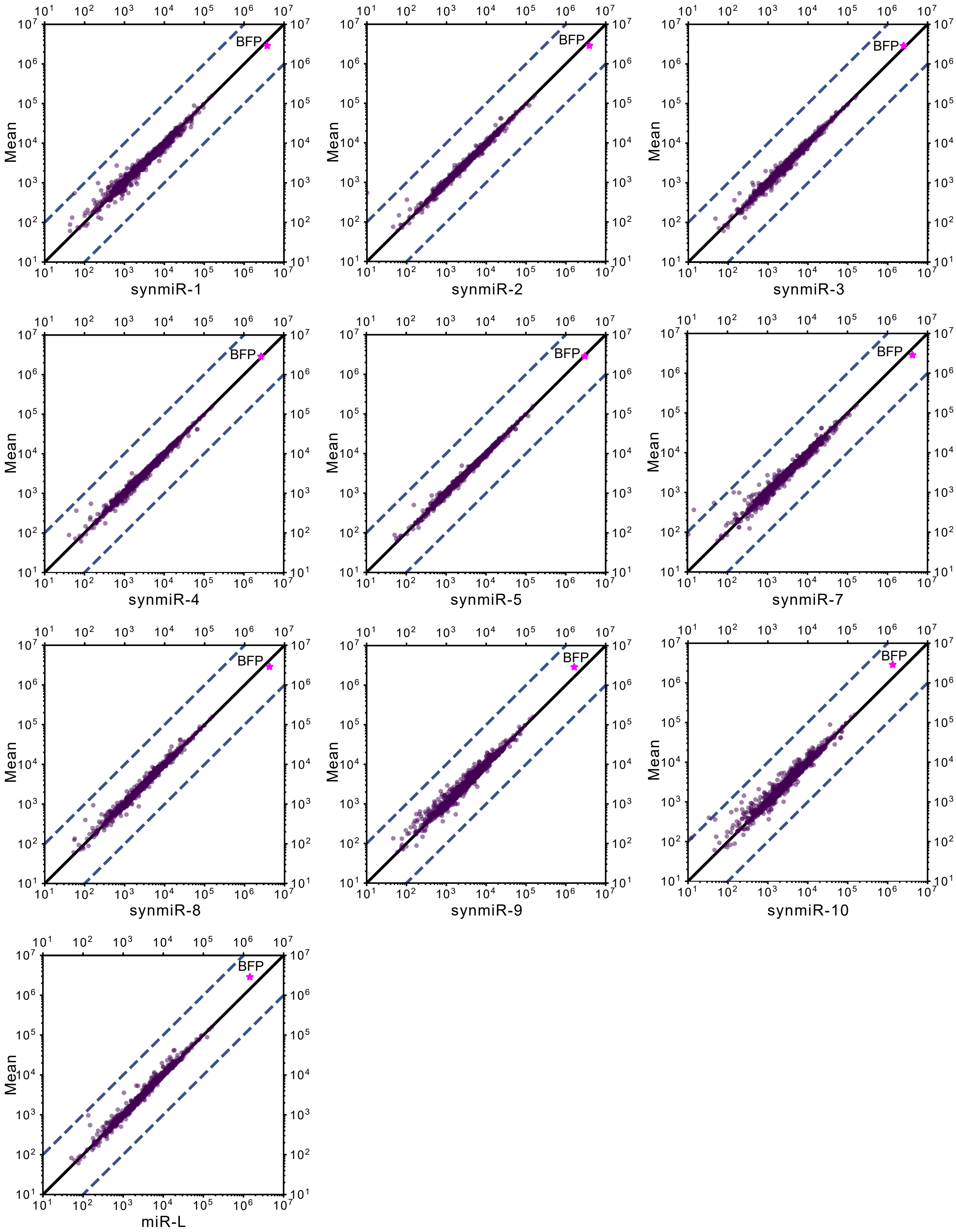


**Data S2-5. Plots of the normalized transcripts per million (TPM) of the synthetic miRNA expressing cells versus the mean TPM, related to Figure 5.**

The mean TPM was calculated by averaging all the TPM of all the synmiR-expressing cell samples. Solid line indicates where the TPM of the sample is equal to the mean. Dashed lines indicate 10 fold expression differences between the mean and the synmiR-expressing cells.

**Data S2-5 Table: Gene Annotation for the significantly differentially expressed genes suggested by bulk RNAseq**.
Shared differentially expressed genes were color-coded.

| miRNA | Differentially expressed genes |
|---|---|
| synmiR-1 | HSPA6, PCSK5, HSPA1A, HSPA1B, KRT17, LOLX4, BAG3, SSC4D, FADS2, GNB2, TUBA1A, TNNC1, CLPTM1L, FOXD1, CCN2 |
| synmiR-2 | HSPA6, PCSK5, HSPA1A, HSPA1B, RPL17 |
| synmiR-3 | HSPA6, PCSK5, HSPA1A, HSPA1B, SPARC |
| synmiR-4 | HSPA6, PCSK5, HSPA1A, HSPA1B, CLPTM1L, POLR2L |
| synmiR-5 | HSPA6, PCSK5, HSPA1A, HSPA1B, CLPTM1L, SSC4D, RPL17, TNNC1 |
| synmiR-7 | HSPA6, PCSK5, HSPA1A, HSPA1B, BAG3, GNB2, AP2S1, CLPTM1L, RHOC, HDLBP |
| synmiR-8 | HSPA6, PCSK5, HSPA1A, HSPA1B, RPL17, LOXL4, FADS2, KRT17, RPS2, SSC4D, SPARC, DDX5, TNNC1 |
| synmiR-9 | HSPA6, LOXL4, FADS2, KRT17, PCSK5, RPL17, HSPA1B, SSC4D, DKK3, LOXL1, TPM1, TUBA1A, BCAM, SPARC, MAGED1, SEMA6B, TNNC1, AP2S1, RPS2, RPS19 |
| synmiR-10 | HSPA6, LOXL4, FADS2, KRT17, PCSK5, GNAS, RPL17, SSC4D, DKK3, BCAM, MAGED1, DDX5, SPARC, TUBA1A, YWHAZ, TNNC1, AP2S1, ACTG1, MRFAP1, RPS2, VIM |
| miR-L | HSPA6, GNAS, HSPA1B, RPL17, VIM |

Data S2-6

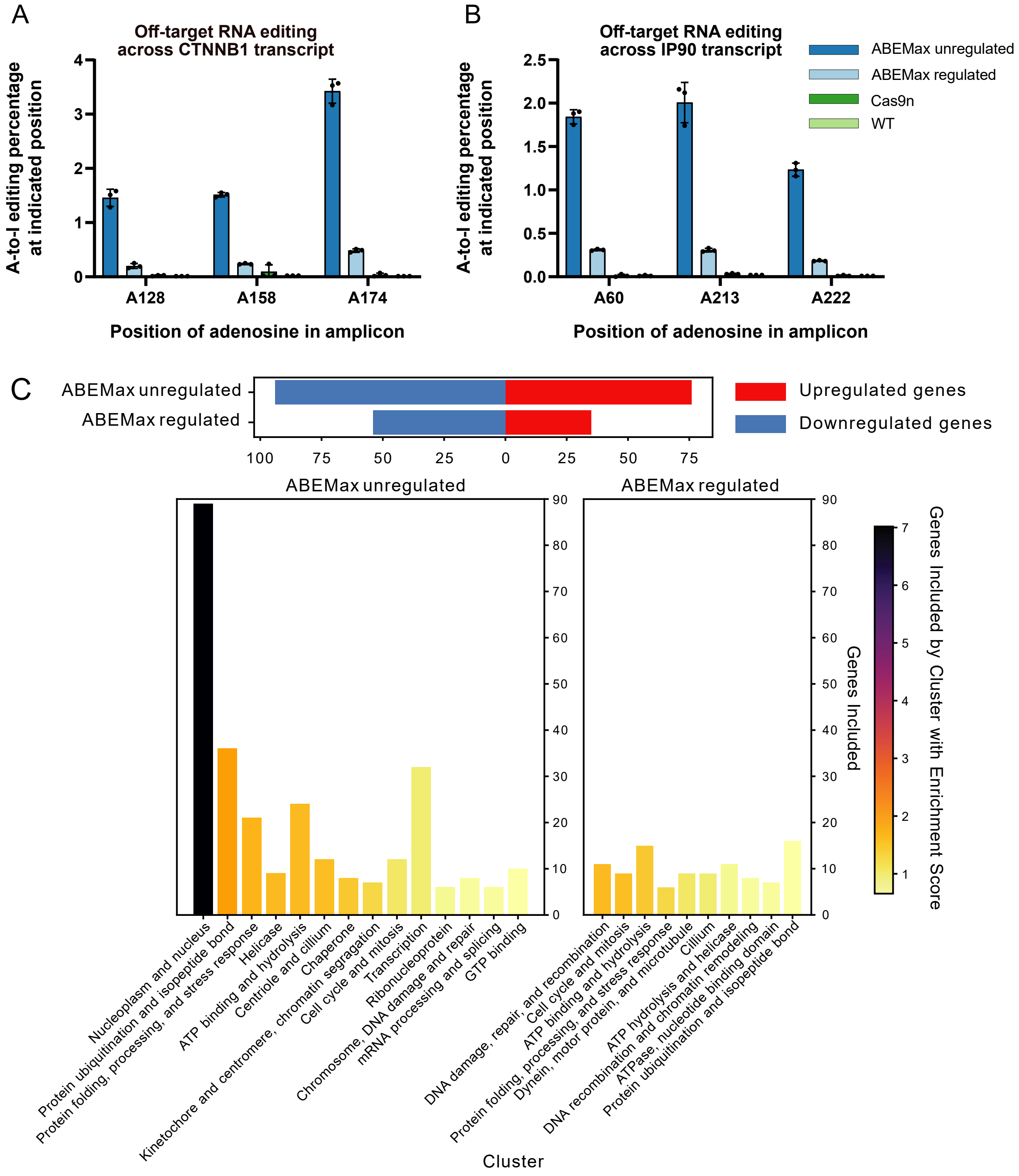


**Data S2-6. DIMMER circuits reduce the off-target RNA editing of the ABEMax base editor, related to Figure 7.** The off-target A-to-I RNA editing percentage of the four specific A sites in the CTNNB1 **(A)** and the IP90 **(B)** transcript. Each dot is a biological replicate. Error bars show the standard deviation.

**(C)** Gene ontology annotation of significantly affected genes identified by bulk RNA sequencing (as in **Figure 7F** and **STAR Methods**). Upper panel, the number of significantly affected genes in the unregulated ABEMax and 4×18nt circuit-regulated ABEMax groups. Lower panel, gene ontology annotation of significantly upregulated/downregulated genes in the unregulated ABEMax group (left) and the 4×18nt circuit-regulated ABEMax group (right). Color bar denotes enrichment score.

**Table S3 DNA sequence of docking strands and imager strands, related to DNA-PAINT in STAR★METHODS.**

| Sequence name | Docking strand sequence (5' to 3') | Imager strand sequence (5' to 3') |
|---|---|---|
| R2 | ACCACCACCACCACCACCA | TGGTGGT-Cy3B |
| R3 | CTCTCTCTCTCTCTCTCTC | GAGAGAG-Cy3B |

**Table S4 The viral titers used in the mice cranial window study, related to DIMMER circuit dynamical quantitation in mice cranial window in STAR★METHODS**. The unit of the viral titers is viral genome copies per mL.

| Group | Cargo | Viral Titers |
|---|---|---|
| CaMKIIα_unregulated | CAP-B10.CaMKIIα.H2B.EGFP.sv40pA<br>CAP-B10.CaMKIIα.H2B.mRuby3.sv40pA | $1.06\times10^{14}$<br>$2.12\times10^{14}$ |
| CaMKIIα_regulated | CAP-B10.CaMKIIα.H2B.EGFP(4×19nt regulated).sv40pA<br>CAP-B10.CaMKIIα.H2B.mRuby3.sv40pA | $8.11\times10^{13}$<br>$2.12\times10^{14}$ |
| MeP_unregulated | CAP-B22.CMVenhancer.MEP229.H2B.EGFP.sv40pA<br><br>CAP-B22.CMVenhancer.MEP229.H2B.mRuby3.sv40pA | $1.22\times10^{14}$<br>$4.04\times10^{13}$<br>$1.94\times10^{14}$ |
| MeP_regulated | CAP-B22.CMVenhancer.MEP229.H2B.EGFP(4×19nt regulated).sv40pA<br>CAP-B22.CMVenhancer.MEP229.H2B.mRuby3.sv40pA | $4.14\times10^{14}$<br>$9.86\times10^{13}$<br>$1.94\times10^{14}$ |
| mDLX2_unregulated | CAP-B10.mDLX2enhancer.H2B.EGFP.sv40pA<br>CAP-B10.mDLX2enhancer.H2B.mRuby3.sv40pA | $1.53\times10^{14}$<br>$1.64\times10^{14}$ |
| mDLX2_regulated | CAP-B10.mDLX2enhancer.H2B.EGFP(4×19nt regulated).sv40pA<br>CAP-B10.mDLX2enhancer.H2B.mRuby3.sv40pA | $7.79\times10^{13}$<br>$4.98\times10^{13}$<br>$1.64\times10^{14}$ |
